# Binding Site-enhanced Sequence Pretraining and Out-of-cluster Meta-learning Predict Genome-Wide Chemical-Protein Interactions for Dark Proteins

**DOI:** 10.1101/2022.11.15.516682

**Authors:** Tian Cai, Li Xie, Shuo Zhang, Muge Chen, Di He, Amitesh Badkul, Yang Liu, Hari Krishna Namballa, Michael Dorogan, Wayne W. Harding, Cameron Mura, Philip E. Bourne, Lei Xie

## Abstract

Discovering chemical-protein interactions for millions of chemicals across the entire human and pathogen genomes is instrumental for chemical genomics, protein function prediction, drug discovery, and other applications. However, more than 90% of gene families remain dark, i.e., their small molecular ligands are undiscovered due to experimental limitations and human biases. Existing computational approaches typically fail when the unlabeled dark protein of interest differs from those with known ligands or structures. To address this challenge, we developed a deep learning framework PortalCG. PortalCG consists of four novel components: (i) a 3-dimensional ligand binding site enhanced sequence pre-training strategy to represent the whole universe of protein sequences in recognition of evolutionary linkage of ligand binding sites across gene families, (ii) an end-to-end pretraining-fine-tuning strategy to simulate the folding process of protein-ligand interactions and reduce the impact of inaccuracy of predicted structures on function predictions under a sequence-structure-function paradigm, (iii) a new out-of-cluster meta-learning algorithm that extracts and accumulates information learned from predicting ligands of distinct gene families (meta-data) and applies the meta-data to a dark gene family, and (iv) stress model selection that uses different gene families in the test data from those in the training and development data sets to facilitate model deployment in a real-world scenario. In extensive and rigorous benchmark experiments, PortalCG considerably outperformed state-of-the-art techniques of machine learning and protein-ligand docking when applied to dark gene families, and demonstrated its generalization power for off-target predictions and compound screenings under out-of-distribution (OOD) scenarios. Furthermore, in an external validation for the multi-target compound screening, the performance of PortalCG surpassed the human design. Our results also suggested that a differentiable sequence-structure-function deep learning framework where protein structure information serve as an intermediate layer could be superior to conventional methodology where the use of predicted protein structures for predicting protein functions from sequences. We applied PortalCG to two case studies to exemplify its potential in drug discovery: designing selective dual-antagonists of Dopamine receptors for the treatment of Opioid Use Disorder, and illuminating the undruggable human genome for targeting diseases that do not have effective and safe therapeutics. Our results suggested that PortalCG is a viable solution to the OOD problem in exploring the understudied protein functional space.

**Author Summary:** Many complex diseases such as Alzheimer’s disease, mental disorders, and substance use disorders do not have effective and safe therapeutics due to the polygenic nature of diseases and the lack of thoroughly validate drug targets and their ligands. Identifying small molecule ligands for all proteins encoded in the human genome will provide new opportunity for drug discovery of currently untreatable diseases. However, the small molecule ligand of more than 90% gene families is completely unknown. Existing protein-ligand docking and machine learning methods often fail when the protein of interest is dissimilar to those with known functions or structures. We develop a new deep learning framework PortalCG for efficiently and accurately predicting ligands of understudied proteins which are out of reach of existing methods. Our method achieves unprecedented accuracy over state-of-the-arts by incorporating ligand binding site information and sequence-to-structure-to-function paradigm into a novel deep meta-learning algorithms. In a case study, the performance of PortalCG surpassed the human design. The proposed computational framework will shed new light into how chemicals modulate biological system as demonstrated by applications to drug repurposing and designing polypharmacology. It will open a new door to developing effective and safe therapeutics for currently incurable diseases. PortalCG can be extended to other scientific inquiries such as predicting protein-protein interactions and protein-nucleic acid recognition.

## 1 Introduction

The central aim of scientific inquiry has been to deduce new concepts from existing knowledge or to generalize observations. Numerous such issues exist in the biological sciences. The rise of deep learning has sparked a surge of interest in using machine learning to explore previously unexplored molecular and functional spaces in biology and medicine, ranging from “deorphanizing” G-protein coupled receptors[1] and translating cell-line screens to patient drug responses[2][3], to predicting novel protein structures[4][5][6], to identifying new cell types from single-cell omics data[7]. Illuminating the understudied space of human knowledge is a fundamental problem that one can attempt to address via deep learning—that is, to generalize a “well-trained” model to unseen data that lies Out-of-Distribution (OOD) of the training data, in order to successfully predict outcomes under conditions that the model has never encountered before. While deep learning is capable, in theory, of simulating any function mapping, its generalization power is notoriously limited in the case of distribution shifts[8].

The training of a deep learning model starts with a domain-specific model architecture. The final model instance selected and its performance are determined by a series of data-dependent design choices, including model initialization, data used for training/validation/testing, optimization of loss function, and evaluation metrics. Each of these design choices impacts the generalization power of a trained model. The development of several recent deep learning-based approaches—notably transfer learning[9], self-supervised representation learning[10], and meta-learning[11][12]—has been motivated by the OOD challenge. However, each of these methods focuses on only one aspect in the training pipeline of a deep neural network model. Causal learning and mechanism-based modeling (e.g., based on the first principle of physics) could be a more effective solution to the OOD problem [8], but at present these approaches can be applied only on modest scales because of data scarcity, computational complexity, or limited domain knowledge. Solving large-scale OOD problems in biomedicine via machine learning would benefit from a systematic framework for integrative, beginning-to-end model development and deployment and the incorporation of domain knowledge into the training process.

Here, we propose a new deep learning framework, Portal learning of Chemical Genomics (PortalCG) for predicting small-molecule binding to dark proteins whose ligands are unknown and dark gene families in which all protein members do not have known ligands. Here Portal represents multiple training components in an end-to-end deep learning framework used to systematically address OOD challenges. Small molecules act as endogenous or exogenous ligands of proteins, assisting in maintaining homeostasis of a biological system or severing as therapeutics agents to alter pathological processes. Despite tremendous progress in high-throughput screening, the majority of chemical genomics space remains unexplored[13] due to high costs, inherent limitations in experimental approaches, and human biases[14][15]. Even in well-studied gene families such as G-protein coupled receptors (GPCRs), protein kinases, ion channels, and estrogen receptors, a large portion of proteins remain dark[13]. Elucidating dark proteins and gene families can shed light on many essential but poorly understood biological processes, such as microbiome-host interactions mediated by metabolite-protein interactions. Such efforts could also be instrumental for drug discovery. Firstly, although conventional one-drug-one-gene drug discovery process intends to screen drugs against a single target, unrecognized off-target effects are a common occurrence[16]. The off-target is either the cause of undesirable side effects or present unique potential for drug repurposing. Secondly, polypharmacology, i.e., designing drugs that can target multiple proteins, is needed to achieve desired therapeutic efficacy and combat drug resistance for multi-genic diseases[16]. Finally, identifying new druggable targets and discovering their ligands may provide effective therapeutic strategies for currently incurable diseases; for instance, in Alzheimer’s disease (AD), many disease-associated genes have been identified through multiple omics studies, but are presently considered undruggable[17].

Accurate and robust prediction of chemical-protein interactions (CPIs) across the genome is a challenging OOD problem[1]. If one considers only the reported area under the receiver operating characteristic curve (AUROC), which has achieved 0.9 in many state-of-the-art methods[18][19], it may seem the problem has been solved. However, existing methods have rarely been applied to dark gene families. The performance has been primarily measured in scenarios where the data distribution in the test set does not differ significantly from that in the training set, in terms of similarities between proteins or between chemicals. Few sequence-based methods have been developed and evaluated for an out-of-gene family scenario, where proteins in the test set belong to different (non-homologous) gene families from those in the training set; this sampling bias is even more severe in considering cases where the new gene family does not have any reliable three-dimensional (3D) structural information. Therefore, one can fairly claim that all existing work has been confined to just narrow regions of chemical genomics space for an imputation task, without validated generalizability into the dark proteins for novel discoveries. We have shown that PortalCG significantly outperforms the leading machine learning and protein-ligand docking methods that are available for predicting ligand binding to dark proteins. Thus, PortalCG may shed new light on unknown functions for dark proteins, and empower drug discovery using Artificial Intelligence (AI). To demonstrate the potential of PortalCG, we applied PortalCG to two case studies: designing selective dual-antagonists of Dopamine receptors for Opioid Use Disorder (OUD) with experimental validations, and illuminating the understudied druggable genome for targeting diseases that lack effective and safe therapeutics. The novel genes and their lead compounds identified from PortalCG provide new opportunities for drug discovery to treat currently incurable diseases such as OUDs, and Alzheimer’s disease (AD). They warrant further experimental validations.

In summary, the contributions of this work are two-fold:

1. We proposed a new algorithm PortalCG to improve the generalization power of machine learning on OOD problems. Comprehensive benchmark studies demonstrate the promise of PortalCG when applied to exploring the dark gene families in which proteins do not have any known small molecule ligands.
2. Using PortalCG, we shed new light on unknown protein functions in dark proteins (viz. small molecule-binding properties), and open new avenues in polypharmacology and drug repurposing; as demonstrated by identifying novel drug targets and lead compounds for OUDs and AD.

## 2 Results and Discussion

### 2.1 Overview of PortalCG

PortalCG includes four key biology-inspired components: 3-dimensional (3D) binding site-enhanced sequence pre-training, end-to-end sequence-structure-function step-wise transfer learning (STL), out-of-cluster meta-learning (OOC-ML), and stress model selection (see Figure 1).

**Figure 1:**
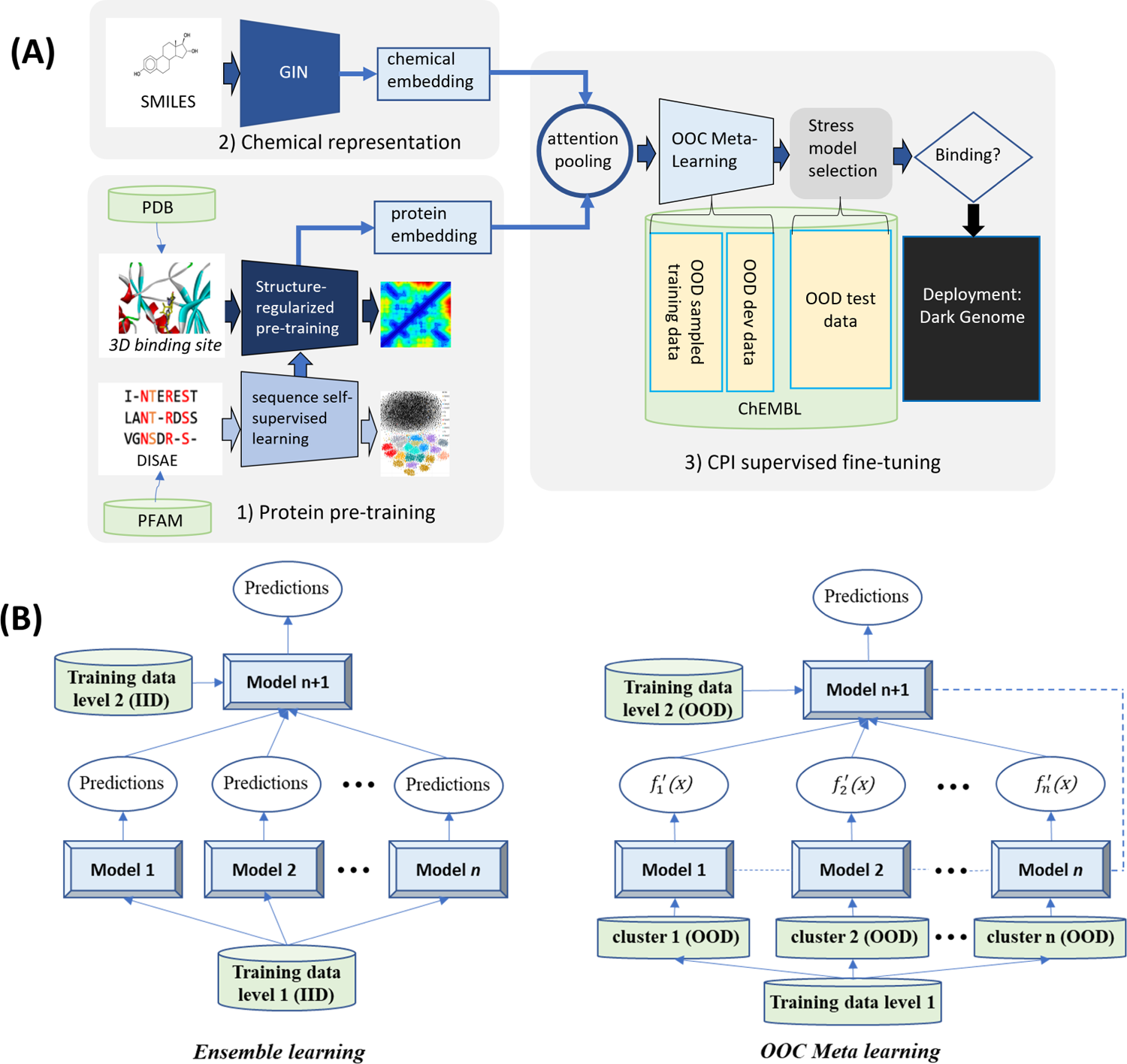
**(A) Scheme of PortalCG**: PortalGC enables chemical protein interactions (CPIs) prediction for dark proteins across gene families with four key components: ligand binding site enhanced sequence pretraining, end-to-end transfer learning following sequence-structure-function paradigm, out-of-cluster meta-learning (OOC-ML), and stress model selection. **(B) Illustration of OOC-ML with the comparision with classic ensemble learning**: OOC-ML follows the same spirit as the ensemble learning, but different in data split, model architecture, and optimization schema.

#### 3D binding site-enhanced sequence pre-training

Pre-training strategy is a proven powerful approach to boost the generalizability of deep learning models[20]. Pre-trained natural language models have revolutionized Natural Language Processing (NLP)[20]. Significant improvements are also observed when applying the same pre-training strategy to protein sequences for structure[5], function[21][22], and CPI predictions[1]. We begin by performing self-supervised training to map tens of millions of sequences into a universal embedding space, using state-of-the-art *distilled sequence alignment embedding* (DISAE) algorithm [1]. In brief, DISAE first distills the original sequence into an ordered list of amino acid triplets by extracting evolutionarily important positions from a multiple sequence alignment. Then long-range residue-residue interactions can be learned via the Transformer module in ALBERT[10]. A self-supervised masked language modeling (MLM) approach is used where 15% triplets are randomly masked and assumed as unknown and the remaining triplets are used to predict what the masked triplets are. In this way, DISAE leaned the protein sequence representation to capture functional information without the knowledge of their structure and function.

3D structural information about the ligand-binding site was used to fine-tune the sequence embedding because it can be evolutionarily related across the fold space and is more informative than the sequence alone for the ligand binding[23]. On the top of pre-trained DISAE embeddings, amino acid residue-ligand atom distance matrices generated from protein-ligand complex structures were predicted from the protein sequence and its ligand. As a result, the original DISAE embedding was re-fined by the 3D ligand binding site information. This structure-regularized protein embedding was used as a hidden layer for supervised learning of cross-gene family CPIs, following an end-to-end sequence-structure-function training process described below.

#### End-to-end sequence-structure-function STL

The function of a protein—e.g., serving as a target receptor for ligand binding—stems from its three-dimensional (3D) shape and dynamics which, in turn, is ultimately encoded in its primary amino acid sequence. In general, information about a protein’s structure is more powerful than purely sequence-based information for predicting its molecular function because sequences drift/diverge far more rapidly than do 3D structures on evolutionary timescales. Furthermore, proteins from different gene families may have similar functional sites, thus perform similar functions[23]. Although the number of experimentally-determined structures continues to exponentially increase, and now AlphaFold2 can reliably predict 3D structures of many single-domain proteins, it nevertheless remains quite challenging to directly use protein structures as input for predicting ligand-binding properties of dark proteins. This motivates us to directly use protein sequences to predict ligands of dark proteins in PortalCG. Protein structure information is used as an intermediate layer as trained by the structure-enhanced pre-training to connect a protein sequence and a corresponding protein function (Figure 1A), as inspired by the concept of differentiable biology[24]. By encapsulating the role of structure in this way, inaccuracies and uncertainties in structure prediction are “insulated” and will not propagate to the function prediction. Details of neural network architecture and training methods can be found in section 4.2.

#### Out-of-cluster meta-learning (OOC-ML)

We designed a new OOC-ML approach to explore dark gene families. Here, predicting ligands of dark gene families can be formulated as the following problem: how can we quickly learn the ligand binding pattern of a new gene family without labeled data from the information obtained from other gene families with a relatively large amount of labeled data? Meta-learning is a general learning strategy that learns a new task without or with few labeled data from outputs (meta-data) generated by multiple other tasks with labeled data, thus naturally fits our purpose. The principle of OOC-ML is first to independently learn the pattern of ligand bindings from each gene family that has labeled data and then to extract the common intrinsic pattern shared by these gene families and apply the learned essential knowledge to dark ones. OOC-ML is similar to ensemble learning that uses a machine learning model at the high level (the second level) to learn how to best combine the predictions from other machine learning models at the low level (the first level), as shown in Figure1B. Nevertheless, there are three key differences between proposed OOC-ML and classic ensemble learning. First, all low-level models in the ensemble learning use the same training data, and the training data used in the high-level has the same distribution as that used in the low-level. In the OOC-ML, the training data for each low-level model has a different distribution. Specifically, they come from different Pfam families. The training data in the high-level also uses Pfam families that are different from all others used in the low-level. Second, instead of using different machine learning algorithms in the low-level ensemble model, the model architecture for all models in the OOC meta-learning is the same as inspired by Model Agnostic Meta-Learning (MAML)[11]. The difference between models lies in their different parameters (mapping functions) due to the different input data. Finally, ensemble learning uses the predictions from the low-level models as meta data for the input of the high-level model. OOC meta-learning uses gradients of mapping functions of the low-level models as meta data, which represent how the model learns, and retrains the gradients by the high-level model.

#### Stress model selection

Finally, training should be stopped at a suitable point in order to avoid overfitting. This was achieved by stress model selection. Stress model selection is designed to basically recapitulate an OOD scenario by splitting the data into OOD train, OOD development, and OOD test sets as listed in Table 1; in this procedure, the data distribution for the development set differs from that of the training data, and the distribution of the test data set differs from both the training and development data. The section 4.2 provides further methodological details, covering data pre-processing, the core algorithm, model configuration, and implementation details.

**Table 1:** Data split for stress model instance selection

| data split | Common practice | classic scheme applied in OOD | PortalCG | specification |
| --- | --- | --- | --- | --- |
| train | IID train | IID train | / | each batch includes data from the same gene family |
|  | / | / | OOD train | data from different gene families are used in each batch |
| dev | IID-dev | IID-dev | / | from the same gene family as that in the train set |
|  | / | / | OOD-dev | from a different gene family from the training set |
| test | IID-test | / | / | from the same gene family as that in the training set |
|  | / | OOD-test | OOD-test | from a different gene family from both OOD-dev and training set |

### 2.2 There are significantly unexplored dark gene families for small molecule binding

We inspected the known CPIs between (i) molecules in the manually-curated ChEMBL database, which consists of only a small portion of the chemical space, and (ii) proteins annotated in Pfam-A [25], which represents only a narrow slice of the whole protein sequence space. The ChEMBL26[26] database supplies 1, 950, 765 chemicals paired to 13, 377 protein targets, constituting 15, 996, 368 known interaction pairs. Even for just this small portion of chemical genomics space, unexplored gene families are enormous, can be seen in the dark region in Figure 2. Approximately 90% of Pfam-A families do not have any known small-molecule binder. Even in Pfam families with annotated CPIs (e.g., GPCRs), there exists a significant number of “orphan” receptors with unknown cognate ligands (Figure 2). Fewer than 1% of chemicals bind to more than two proteins, and < 0.4% of chemicals bind to more than five proteins, as shown in Supplemental Figure S1, S2 and S3. Because protein sequences in the dark gene families could be significantly different from those for the known CPIs, predicting CPIs for dark proteins is an archetypal, unaddressed OOD problem.

**Figure 2:**
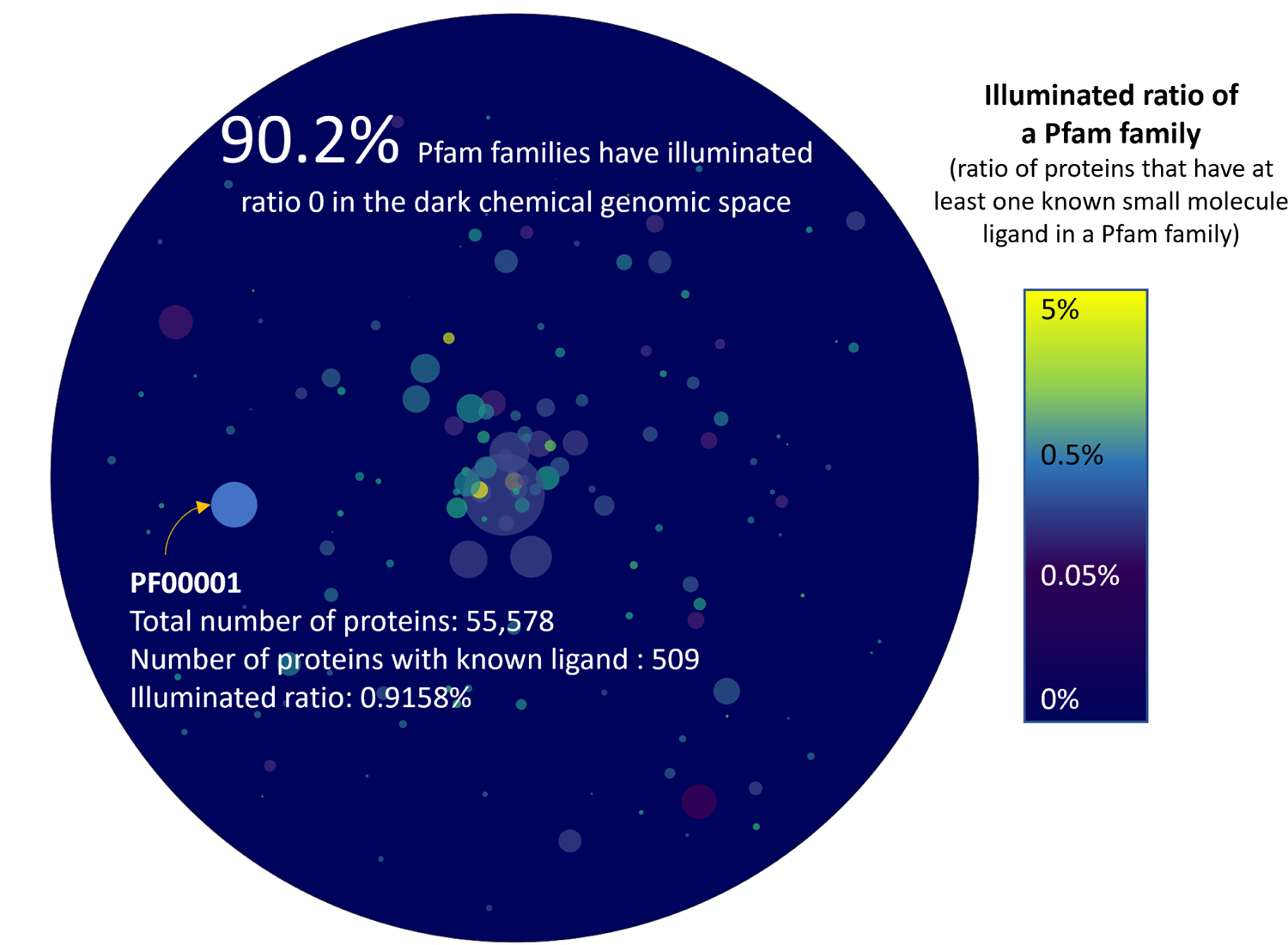
Chemical genomics space in statistics: The ratio of proteins that have at least a known ligand in each Pfam family. Each color bubble represents a Pfam family. The size of a bubble is proportional to the total number of proteins in the Pfam family. 1, 734 Pfam families have at least one known small molecule ligand. Most of these Pfam families have less than 1% proteins with known ligands. Furthermore, around 90.2% of total 17, 772 Pfam families remain dark without any known ligand information.

### 2.3 PortalCG significantly outperforms state-of-the-art approaches to predicting CPIs of dark gene families

Two major categories of approaches have been developed for CPI predictions: machine learning and protein-ligand docking (PLD). Recently published DISAE has been shown to outperform other leading deep learning methods for predicting CPIs of orphan receptors and is explainable[1]. Because the neural network architecture of PortalCG is similar to that of DISAE, we used DISAE as the baseline to evaluate the performance improvement of PortalCG over the state-of-the-art. PortalCG demonstrates superior performance in terms of both Receiver Operating Characteristic (ROC) and Precision-Recall (PR) curves when compared with DISAE, as shown in Figure 3(A). When the ratio of positive and negative cases is imbalanced, the PR curve is more informative than the ROC curve. The PR-AUC of PortalCG and DISAE is 0.714 and 0.603, respectively. In this regard, the performance gain of PortalCG (18.4%) is significant (p-value < 1*e* – 40). Performance breakdowns for binding and non-binding classes can be found in Supplemental Figure S4. PortalCG exhibits much higher recall and precision scores for positive cases (i.e., a chemical-protein pair that is predicted to bind) versus negative, as shown in Supplemental Figure S4; this is a highly encouraging result, given that there are many more negative (non-binding) than positive cases. The deployment gap, shown in Figure 3(B), is steadily around zero for PortalCG; this promising finding means that we can expect that, when applied to the dark proteins, the performance will be similar to that measured using the development data set.

**Figure 3:**
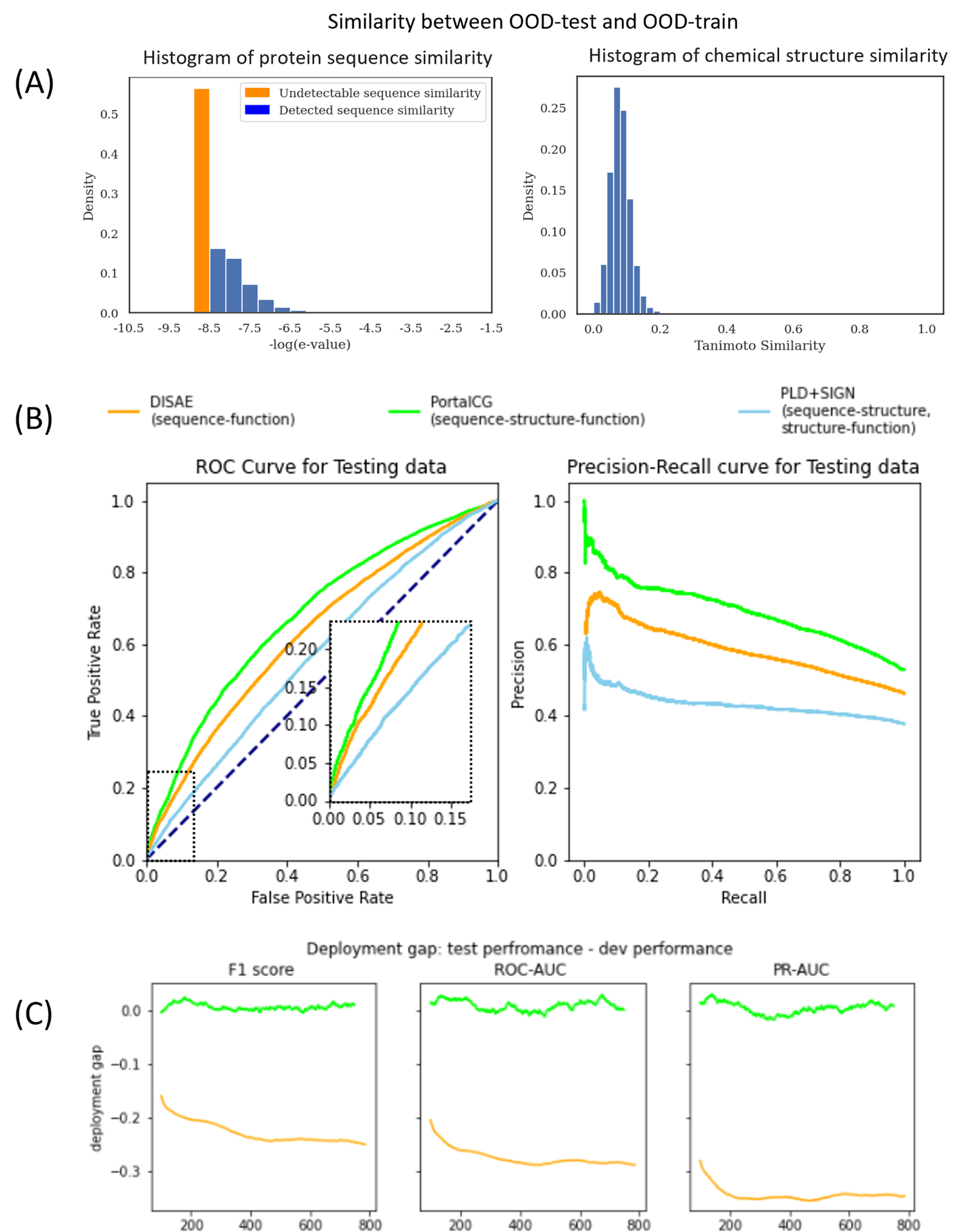
Comparison of PortalCG with the state-of-the-art methods DISAE and PLD+SIGN as baselines using the OOD test where the proteins in the testing data come from different Pfam families from the proteins in the training and validation data. (A) Histograms of protein sequence and chemical structure similarities between OOD-train and OOD-test. The majority of protein sequences in the training set does not have detectable similarity to the proteins in the testing set. (B) Receiver Operating Characteristic (ROC) and Precision-Recall curves for the “best” model instance selected by the stress test. Due to the imbalanced active/inactive data, the Precision-Recall (PR) curve is a more reliable measure than the ROC curve. (C) Deployment gaps of PoralCG and DISAE. The deployment gap of PortalCG is steadily around zero as training step increases while the deployment performance of DISAE deteriorates.

With the advent of high-accuracy protein structural models, predicted by AlphaFold2 [5], it now becomes feasible to use reversed protein-ligand docking (PLD) [27] to predict ligand-binding sites and poses on dark proteins, on a genome-wide scale. In order to compare our method with the reversed protein-ligand docking approach, blind PLD to proteins in the benchmark was performed via Autodock Vina[28] followed by protein-ligand binding affinity prediction using a leading graph neural network-based method SIGN [29] (PLD+SIGN). The binding affinities predicted by SIGN was more accurate than original scores from Autodock Vina (Supplemental Figure S5). The performance of PLD+SIGN was compared with that of PortalGC and DISAE. As shown in Figure 3(A), both ROC and PR for PLD+SIGN are significantly worse than for PortalGC and DISAE. It is well known that PLD suffers from a high false-positive rate due to poor modeling of protein dynamics, solvation effects, crystallized waters, and other challenges [30]; often, small-molecule ligands will indiscriminately “stick” to concave, pocket-like patches on protein surfaces. For these reasons, although AlphaFold2 can accurately predict many protein structures, the relatively low reliability of PLD still poses a significant limitation, even with a limitless supply of predicted structures [31]. Thus, the direct application of PLD remains a challenge for predicting ligand binding to dark proteins. PortalCG’s end-to-end sequence-structure-function learning could be a more effective strategy in terms of both accuracy and efficacy: protein structure information is not used as a fixed input, but rather as an intermediate layer that can be tuned using various structural and functional information. Furthermore, the inference time of PortalCG for predicting a CPI is several orders of magnitude faster than that needed by the PLD. For example, it takes approximate 1 millisecond for PortalCG to predict a ligand binding to DRD2, while Autodock Vina needs around 10 seconds to dock a ligand to DRD2 excluding the time for defining the binding pocket.

### 2.4 Both the STL and OOC-ML stages contribute to the improved performance of PortalCG

To gauge the potential contribution of each component of PortalCG to the overall system effectiveness in predicting CPIs for dark proteins, we systematically compared the four models shown in Table 2. Details of the exact model configurations for these experiments can be found in the Supplemental Materials Table S1. As shown in Table 2, Variant 1, with a higher PR-AUC compared to the DISAE baseline, is the direct gain from transfer learning through 3D binding site information, all else being equal; yet, with transfer learning alone and without OOC-ML as an optimization algorithm in the protein function CPI prediction (i.e., Variant 2 versus Variant 1), the PR-AUC gain is minor. Variant 2 yields a 15% improvement while Variant 1 achieves only a 4% improvement over DISAE. PortalCG, in comparison, has the best PR-AUC score. With all other factors held constant, the advantage of PortalCG appears to be the synergistic effect of both STL and OOC-ML. The performance gain measured by PR-AUC under a shifted evaluation setting is significant (p-value < 1e-40), as shown in Supplemental Figure S6.

**Table 2:** Ablation study of PortalCG.

| Models | Configuration | OOD-test set |  | Deployment gap |  |
| --- | --- | --- | --- | --- | --- |
|  |  | ROC-AUC | PR-AUC | ROC-AUC | PR-AUC |
| PortalCG | PortalCG with all components | 0.677±0.010 | 0.714±0.010 | 0.010±0.009 | 0.005±0.010 |
| DISAE | PotalCG w/o STL or OOC-ML | 0.636±0.004 | 0.603±0.005 | -0.275±0.016 | -0.345±0.012 |
| variant 1 | PotalCG w/o OOC-ML | 0.661±0.004 | 0.629±0.005 | / | / |
| variant 2 | PotalCG w/o STL | 0.654±0.062 | 0.698±0.015 | / | / |
| PLD+SIGN | / | 0.569 | 0.433 | / | / |

We find that stress model selection is able to mitigate potential overfitting problems, as expected. Training curves for the stress model selection are in Supplemental Figure S7. As shown in Supplemental Figure S7, the baseline DISAE approach tends to over-fit with training, and IID-dev performances are all higher than PortalCG but deteriorate in OOD-test performance. Hence, the deployment gap for the baseline is −0.275 and −0.345 on ROC-AUC and PR-AUC, respectively, while PortalCG deployment gap is around 0.01 and 0.005, respectively.

### 2.5 PortalCG is competitive on virtual compound screening for novel chemicals

Given that the pretraining, OOC-ML, and stress tests were only applied to proteins, current PortalCG was primarily designed to explore the dark protein space instead of new chemical space. Nevertheless, we evaluated if PortalCG could improve the performance for compound screening for novel chemicals. We employed a widely used DUD-E benchmark that included 8 protein targets along with their active compounds and decoys[32], and compared the performance of PortalCG with that of PLD. We used DUD-E chemicals as testing set. We trained PortalCG by excluding target proteins in the training/validation sets, and have all chemicals in the training/validation set dissimilar to those in the testing set (Tanimoto Coefficient (TC) less than 0.3 or 0.5). Under these chemical similarity thresholds, the false positive rate in the training/validation set was higher than 95.0% assumed that the ratio of actives vs inactives was 1:50 (Supplemental Figure S8).

As shown in Table 3, except targets kif11 and gcr, PortalCG could surprisingly outperform Autodock Vina on other remaining 6 targets in terms of enrichment factors (EFs). Similarly, PortalCG exhibited higher EFs than PLD-SIGN on six proteins. For the EF of 1%, the compound screening performance of PortalCG on 87.5% and 100.0% of targets is better than random guesses (EF=1.0) when the chemical similarity between the queries and the training data is 0.3 and 0.5, respectively. In contrast, only 50.0% 75.0% targets are better than the random guess for Autodock Vina and PLD+SIGN, respectively. It implies that PortalCG has learned certain patterns of CPIs although the chemical OOD has not explicitly modeled. Different from PLD, whose EFs varied greatly across targets, the variance of EFs was relatively small for PortalCG across the targets, suggesting that the model was not biased to certain proteins. Thus, PortalCG is complementary with PLD, and has a potential to improve the capability of virtual compound screening, especially, for dark proteins whose reliable structures are not available.

**Table 3:**
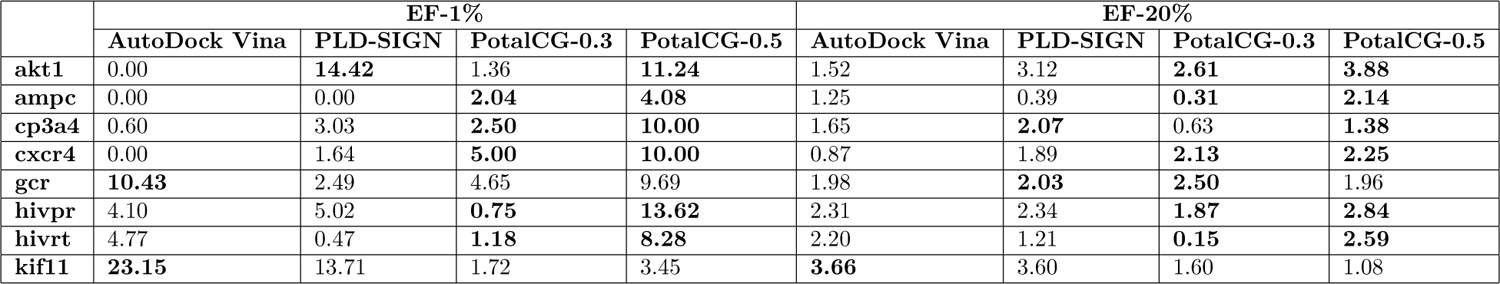
Performances of compound screening evaluated using DUD-E benchmark. PortalCG-0.3: the similarities between chemicals in the training/validation set and those in the testing set are less than 0.3 of Tanimoto Coefficient (TC). PortalCG-0.5: the similarities between chemicals in the training/validation set and those in the testing set are less than 0.5 of TC. The best performance is in bold

### 2.6 PortalCG is capable of screening selective multi-targeted compounds with novel scaffold for dark proteins

Opioid use disorder (OUD) is an overwhelming healthcare and economic burden. Although several pharmaceutical treatments for OUD exist, they are either restricted in usage or limited in effectiveness. Dopamine D1 and D3 receptors (DRD1 and DRD3) have been identified as potential drug targets for OUD. DRD1 partial agonists and antagonists alter the rewarding effects of drugs, while DRD3 antagonists reduce drug incentive and behavioral responses to drug cues. Moreover, recent evidence suggests that simultaneous targeting of DRD1 and DRD3 may be an effective OUD therapeutic strategy as the combination of a DRD1 partial agonist and a DRD3 antagonist reduced cue-induced relapse to heroin in rats[33]. By contrast, dopamine D2 receptor (DRD2) antagonism is associated with cataleptic side effects which limit the use of DRD2 antagonists as OUD therapeutics. Thus, selective DRD1 and DRD3 dual-antagonists could be an effective strategy for OUD treatment[34]. Because there are multiple dopamine receptors (especially, DRD2) that are similar to D1R and D3R, it is challenging to develop a selective dual-antagonist for DDR1 and DRD3. PortalCG may provide new opportunities for OUD polypharmacology.

We synthesized 65 compounds based on the scaffold as shown in Figure 4A, which combines structural features of the DRD1 antagonist (-)-stepholidine with DRD3 antagonist pharmacophore, and determined their binding affinities to DRD1, DRD2, and DRD3, respectively (supplemental table S2). Tens of thousands of possible chemical structures could be derived from different combinations of R1, R2, R3, R4, and linker functional groups, as marked in Figure 4A. We have little knowledge on what is an optimal combination of functional groups for a dual-DRD1/DRD3 antagonist. If we define an acceptable dual-DRD1/DRD3 antagonist as a compound whose binding affinities are less than 100 nM of Ki to both DRD1 and DRD3 but higher than 100 nM of Ki to DRD2, only 10 compounds satisfied this condition (successful rate of 15.4%) among the 65 synthesized compounds. It notes that a safe therapeutics for OUD may need much lower binding affinity than 100 nM of Ki for DRD2. For the 28 DRD1 antagonists with the Ki less than 100 nM, 15 of them had Ki higher than 100 nM for DRD3, corresponding to a two-class successful rate of 46.4%. It suggested that our current knowledge is limited for effectively designing selective dual-DRD1/3 antagonists using existing scaffolds, let alone under a novel scaffold. The question is if we can use computational methods, especially PortalCG, to identify selective dual-DRD1/3 antagonists with a novel scaffold. We performed a rigorous bind test to validate the performance of PortalCG for this purpose. In the evaluation of PortalCG and DISAE, all of chemicals in the training data had different scaffolds from 65 test compounds, i.e., an OOD scenario on the chemical side [35]. Three models were trained with the sequence similarity between DRD1/2/3 and proteins in the training/validation data ranging from 20% to 60%. The performance was measured by the accuracy of a three-label classifier. When the sequence identifies between DRD1/2/3 and the proteins in the training/validation set were less than 40%, PortalCG achieved 20.0% and 50.7% successful rate for the cases where all DRDs and any two of them were predicted correctly, respectively (Figure 4B). The successful rate for the all DRDs was significantly higher the human design. Decreasing the sequence identifies between the proteins in the training/validation set and DRD1/2/3 from 40% to 20% only slightly lower the accuracy of PortalCG, as shown in Figure 4C. The performance drops were not statistically significant (p-value > 0.05). Increasing the sequence identities from 40% to 60% also did not significantly change the accuracy. Thus, PortalCG by design was robust to OOD data.

**Figure 4:**
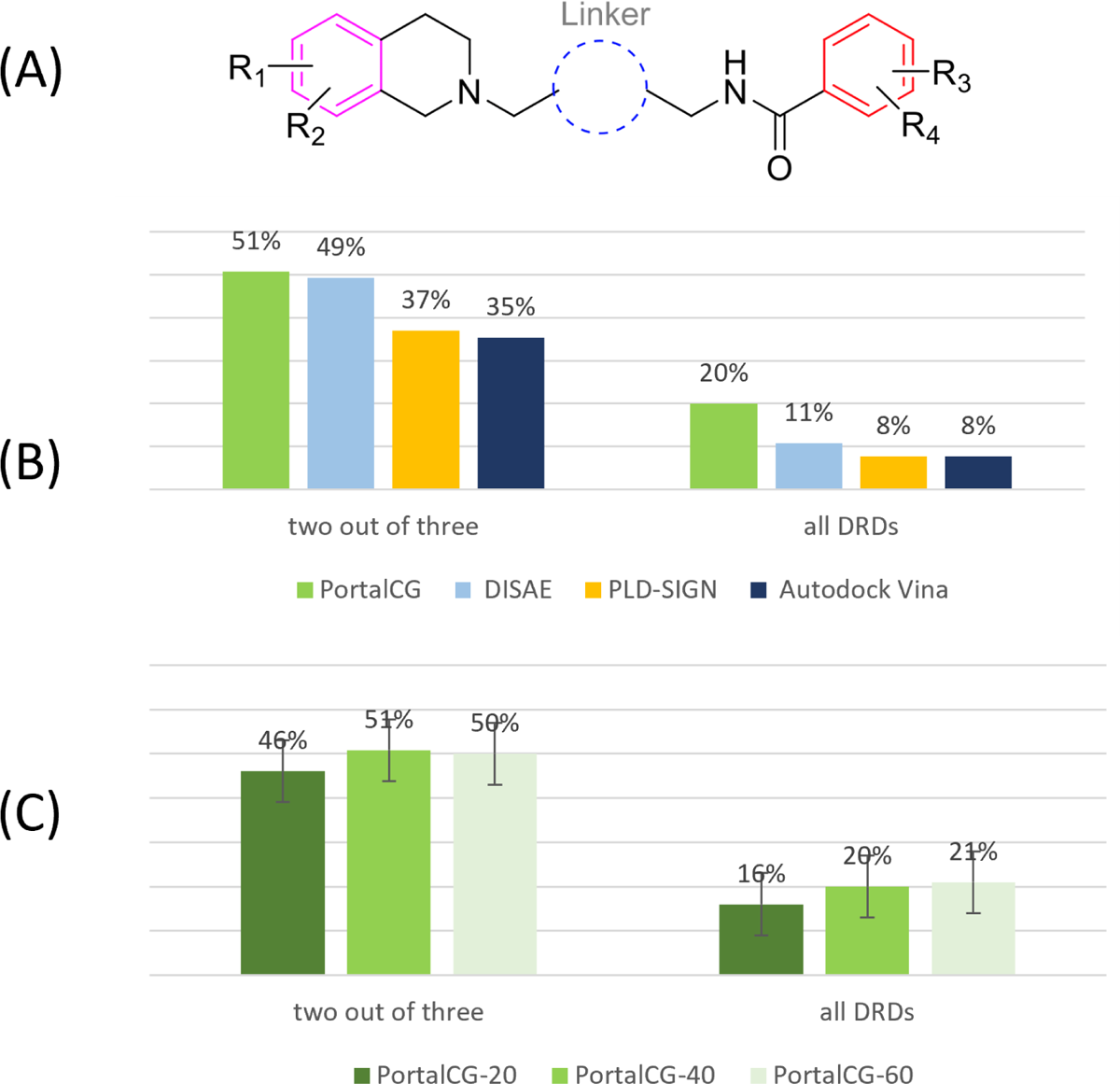
(A) The chemical scaffold on which 65 compounds were synthesized for potential selective dual-DRD1/DRD2 antagonists. Tens of thousands of chemicals can be generated from the different combination of four functional groups R1, R2, R3, and R4 and a linker group. (B) The prediction accuracy of DRD binding profile classification. (C) The performance of PortalCG when the sequence similarities between the proteins in the training/validation set and DRD1/DRD2/DRD3 were less than 20%, 40%, and 60%, respectively. The performance was measured by the accuracy of a three-label classifier. “Two out of three” and “all DRDs” represented the accuracy when two labels and all three labels were predicted correctly.

We compared PortalCG with three baselines DISAE, PLD+SIGN, and Autodock Vina[28]. The crystal structures of DRD1 (PDB id: 7JOZ), DRD2 (PDB id: 6CM4), and DRD3 (PDB id: 3PBL), which were co-crystallized with ligands, were used for the docking. The 65 compounds were docked to the pre-defined binding pocket based on the co-crystallized ligand. The order of accuracy follows PortalCG>DISAE>PLD-SIGN>Autodock Vina, as shown in Figure 4B. This observation is consistent with our benchmark studies. Note that the complex structure was only used for the baseline PLD models but this information was not used for PortalCG and DISAE.

### 2.7 Illuminating the undruggable human genome for drug repurposing

To further demonstrate the potential application of PortalCG, we explored potential drug lead compounds for undrugged disease genes in the dark human genome, and prioritized undrugged genes that can be efficaciously targeted by existing drugs. It is well known that only a small subset of the human genome is considered druggable [36]. Many proteins are deemed “undruggable” because there is no information on their ligand-binding properties or other interactions with small-molecule compounds (be they endogenous or exogenous ligands). Here, we built an “undruggable” human disease protein database by removing the druggable proteins in Pharos [37] and Casas’s druggable proteins [38] from human disease associated genes [17]. A total of 12,475 proteins were included in our disease-associated undruggable human protein list. We applied PortalCG to predict the probability for these “undruggable” proteins to bind to drug-like molecules. Around 6,000 drugs from the Drug Repurposing Hub[39] were used in the screening. The proteins that could bind to a small molecule drug were ranked according to their prediction scores, and 267 of them have a false positive rate lower than 2.18e-05, as listed in the Supplemental Table S3. Table 4 shows the statistically significantly enriched functions of these top ranked proteins as determined by DAVID [40]. The most enriched proteins are involved in alternative splicing of mRNA transcripts. Malfunctions in alternative splicing are linked to many diseases, including several cancers [41][42] and Alzheimer’s disease [43]. However, pharmaceutical modulation of alternative splicing process is a challenging task. Identifying new drug targets and their lead compounds for targeting alternative splicing pathways may open new doors to developing novel therapeutics for complex diseases with few treatment options. In addition, several transcription factors and transcription activity related proteins were identified and listed in Table S4, along with their predicted ligands.

**Table 4:** Functional Annotation enrichment for undruggable human disease associated proteins selected by PortalCG

| David Functional Annotation enrichment analysis |  |  |  |  |
| --- | --- | --- | --- | --- |
| Enriched terms in UniProtKB keywords | Number of proteins involved | Percentage of proteins involved | P-value | Modified Benjamini p-value |
| Alternative splicing | 171 | 66.5 | 7.70E-07 | 2.00E-04 |
| Phosphoprotein | 140 | 54.5 | 2.60E-06 | 3.40E-04 |
| Cytoplasm | 91 | 35.4 | 1.30E-05 | 1.10E-03 |
| Nucleus | 93 | 36.2 | 1.20E-04 | 8.10E-03 |
| Metal-binding | 68 | 26.5 | 4.20E-04 | 2.20E-02 |
| Zinc | 48 | 18.7 | 6.60E-04 | 2.90E-02 |

Diseases associated with these 267 human proteins were also listed in Table 5. Since one protein is always related to multiple diseases, these diseases are ranked by the number of their associated proteins. Most of top ranked diseases are related with cancer development. 21 drugs that are approved or in clinical development are predicted to interact with these proteins as shown in Supplemental Table S5. Several of these drugs are highly promiscuous. For example, AI-10-49, a molecule that disrupts protein-protein interaction between CBFb-SMMHC and tumor suppressor RUNX1, may bind to more than 60 other proteins. The off-target binding profile of these proteins may provide invaluable information on potential side effects and opportunities for drug repurposing and polypharmacology. The drug-target interaction network built for predicted positive proteins associated with Alzheimer’s disease was shown in Figure 5.The target proteins in this network were selected according to the threshold of 0.67. The length of the edges in this network was decided by the prediction scores for these drug-target pairs. The longer the edge is, the lower confidence of the prediction is. Thus if a higher threshold was applied, fewer drug-target pairs will appear in this network. In order to validate the binding activity between the drugs and targets in this network, the PLD was performed between the three most promiscuous drugs, AI-10-49, fenebrutinib, PF-05190457 and their predicted targets. Only those targets with known PDB structures or reliable alpha-fold model structures were used in the docking. Docking scores for the 21 drug-target pairs were listed on Supplemental Table S6. For each of the three drugs, the target with with the lowest docking score (the highest binding affinity) was selected as a representative. Docking conformations and interactions between the drugs and their representative targets were shown in Figure 5. Functional enrichment, disease associations, and top ranked drugs for the undruggable proteins with well-studied biology (classified as Tbio in Pharos) and those excluding Tbio are list in Supplemental Table S7-S11.

**Figure 5:**
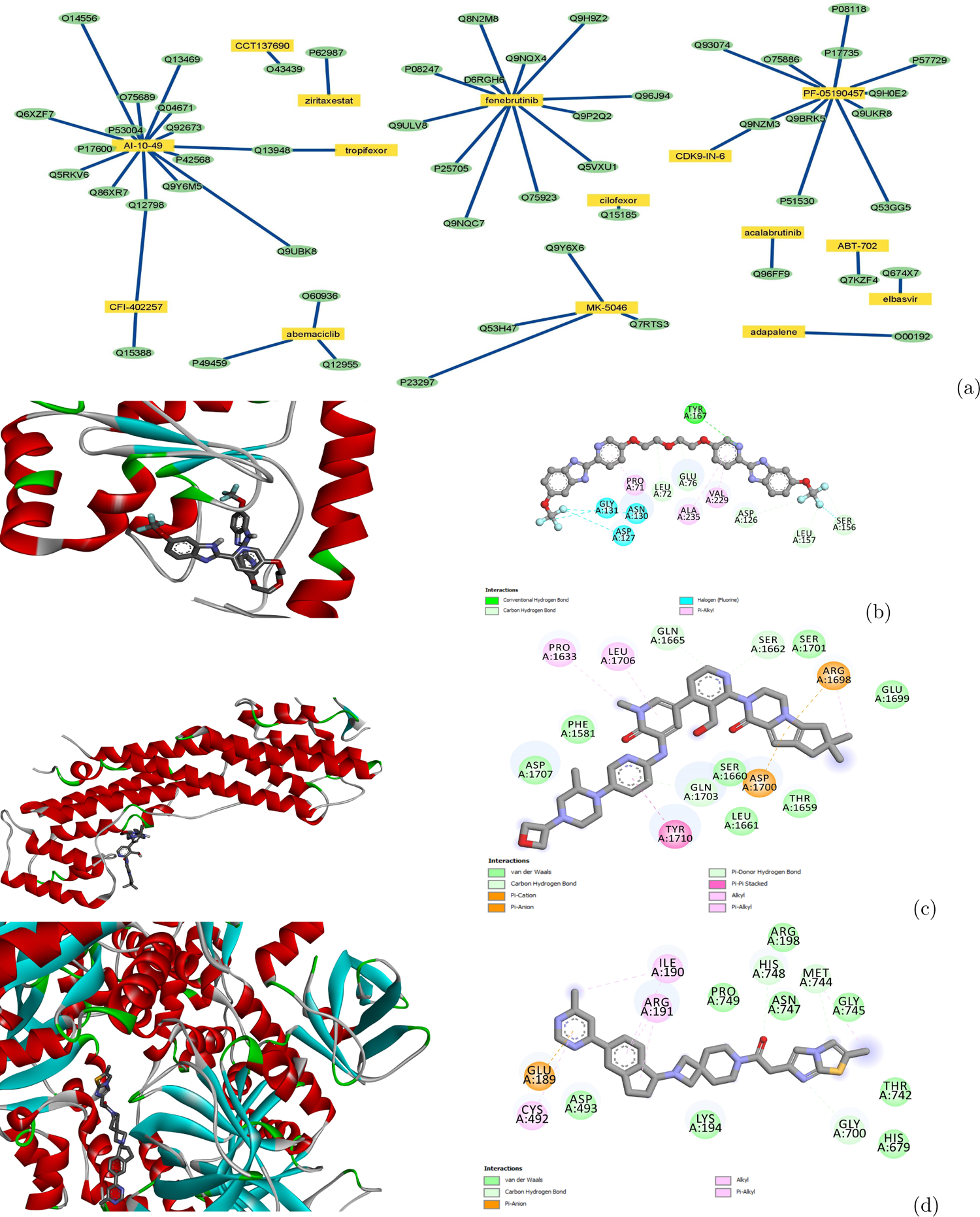
Drug-target interaction network for proteins associated with Alzheimer’s disease and docking poses for representative drug-target pairs calculated by Autodock Vina. (a) Drug-target interaction network predicted by PortalCG. Yellow rectangles and green ovals represent drugs and targets, respectively. (b) Docking pose and ligand binding interactions between protein TIR domain-containing adapter molecule 2 (Uniprot: Q86XR7) and AI-10-49.(c) Docking pose and ligand binding interactions between protein Unconventional myosin-Vc (Uniprot: Q9NQX4) and fenebrutinib. (d) Docking pose and ligand binding interactions between DNA replication ATP-dependent helicase/nuclease (Uniprot: P51530) and PF-05190457.

**Table 5:** Top ranked diseases associated with the undruggable human disease proteins selected by PortalCG

| DiseaseName | # of undruggable proteins associated with disease |
| --- | --- |
| Breast Carcinoma | 90 |
| Tumor Cell Invasion | 86 |
| Carcinogenesis | 83 |
| Neoplasm Metastasis | 75 |
| Colorectal Carcinoma | 73 |
| Liver carcinoma | 66 |
| Malignant neoplasm of lung | 56 |
| Non-Small Cell Lung Carcinoma | 56 |
| Carcinoma of lung | 54 |
| Alzheimer’s Disease | 54 |

## 3 Conclusion

This paper confronts the challenge of exploring dark proteins by recognizing it as an OOD generalization problem in machine learning, and by developing a new deep learning framework to treat this type of problem. We propose PortalCG as a general framework that enables systematic control of the OOD generalization risk. Systematic examination of the PortalCG method revealed its superior performance compared to (i) a state-of-the-art deep learning model (DISAE), and (ii) an AlphaFold2-enabled, GNN-scored, structure-based reverse docking approach. PortalCG showed significant improvements in terms of both sensitivity and specificity, as well as close to zero deployment performance gap. The neural network architecture of PortalCG is similar to DISAE. Its performance improvement over DISAE mainly comes from 3D binding site-enhanced pre-training and OOC-ML optimization. Both PortalCG and DISAE outperform PLD-based methods by getting around the inherent limitations of PLD. Applications of PortalCG to OUD polypharmacology and drug repurposing targeting of hitherto undruggable human proteins affords novel new directions in drug discovery.

PortalCG can be further improved along several directions. In terms of protein sequence modeling, additional *a prior* knowledge of protein structure and function can be incorporated into the pre-training or supervised multi-task learning. The architecture of PortalCG mainly focuses on addressing the OOD problem of protein space but not chemical space. New methods for modeling chemical structures alone or the joint space of chemicals and proteins will no doubt improve CPI predictions for unseen novel chemicals. Future directions include but not limited to novel representation of 3D chemical structures[44] at the sub-molecular level of scaffold and chemical moieties, pre-training of the chemical space [45], and few-shot learning[46] as well as explicitly modeling amino acid-chemical moiety interactions.The existing PortalCG treats the CPI prediction as a binary classification problem, but can be reformulated as a regression model for predicting binding affinities. By defining domain-specific pre-training and down-stream supervised learning tasks, PortalCG could be a general framework to explore the functions of understudied proteins such as protein-protein interactions and protein-nucleic acid recognition.

## 4 Methods

PortalCG as a system level framework involves collaborative new design from data preprocessing, data splitting to model initialization, and model optimization and evaluation. The pipeline of framework is illustrated in Figure 1. Model architecture adopted by PortalCG mostly follows DISAE as shown in Figure 6.

**Figure 6:**
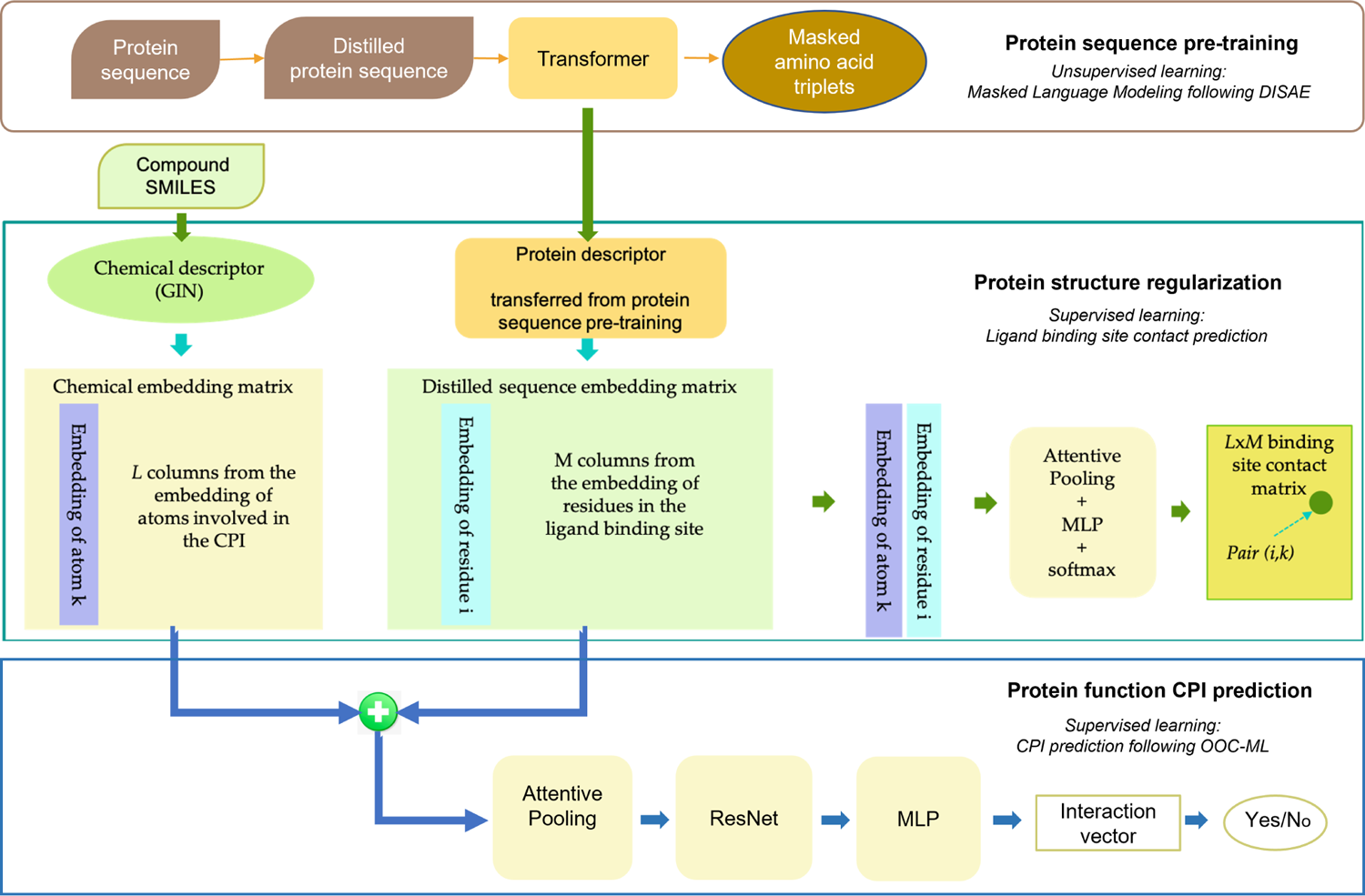
Illustration of PortalCG architecture for three stages of training. The architecture of protein sequence pre-training used a transformer architecture and masked language modeling as detailed in ref[1]. Pretrained protein descriptor was then used in binding site enhanced sequence pre-training. In this stage, the task was to predict amino acid residue and ligand atom distance matrices. Finally, protein descriptors that were pretrained and regularized in the previous two stages were concatenated with chemical descriptors via an attention network to predict CPIs. Chemical structures were represented by GIN[50], a graph neural network model. The second and third stages had same model architecture but the model parameters were transferred from the second to the third stages. OOC-ML as an optimization algorithm was not a model architecture component, and only used in the CPI prediction.

### 4.1 Data sets

PortalCG was trained using four major databases, Pfam[25], Protein Data Bank (PDB)[47], BioLp[48] and ChEMBL[26]. The data were preprocessed as follows.

- Protein sequence data. All sequences from Pfam-A families are used to pretrain the protein descriptor following the same setting in DISAE [1]. DISAE distills the original sequence into an ordered list of amino acid triplets by extracting evolutionarily important positions from a multiple sequence alignment.
- Protein structure. In our protein structure data set, there are 30,593 protein structures, 13,104 ligands, and 91,780 ligand binding sites. Binding sites were selected according to the annotation from BioLip (updated to the end of 2020). Binding sites which contact with DNA/RNA and metal ions were not included. If a protein has more than one ligand, multiple binding pockets were defined for this protein. For each binding pocket, the distances between *Cα* atoms of amino acid residues of the binding pocket were calculated. In order to obtain the distances between the ligand and its surrounding binding site residues, the distances between atom i in the ligand and each atom in the residue j of the binding pocket were calculated and the smallest distance wa selected as the distance between atom i and residue j. In order to get the sequence feature of the binding site residues in the DISAE protein sequence representation[1], binding site residues obtained from PDB structures (queries) were mapped onto the multiple sequence alignments of its corresponding Pfam family. First, a profile HMM database was built for the whole Pfam families. hmmscan [49] was applied to search the query sequence against this profile database to decide which Pfam family it belongs to. For those proteins with multiple domains, more than one Pfam families were identified. Then the query sequence was aligned to the most similar sequence in the corresponding Pfam family by using phmmer. Aligned residues on the query sequence were mapped to the multiple sequence alignments of this family according to the alignment between the query sequence and the most similar sequence.
- Chemical genomics data. CPI classification prediction data is the whole ChEMBL26[26] database where the same threshold for defining positive and negative labels creating as that in DISAE [1] was used. Log-transformation was performed for activities reported in pK_d, pK_i or pIC_50. The activities on a log-scale were then binarized where protein-ligand pairs were considered active if pIC_50> 5.3, pK_d >7.3 or pK_i>7.3.

Above data are split into training, validation, and testing sets. Data split statistics are shown in Table 6. Other data statistics are demonstrated in Figure 2 and Supplemental Materials Figure S1, S2, S3.

**Table 6:**
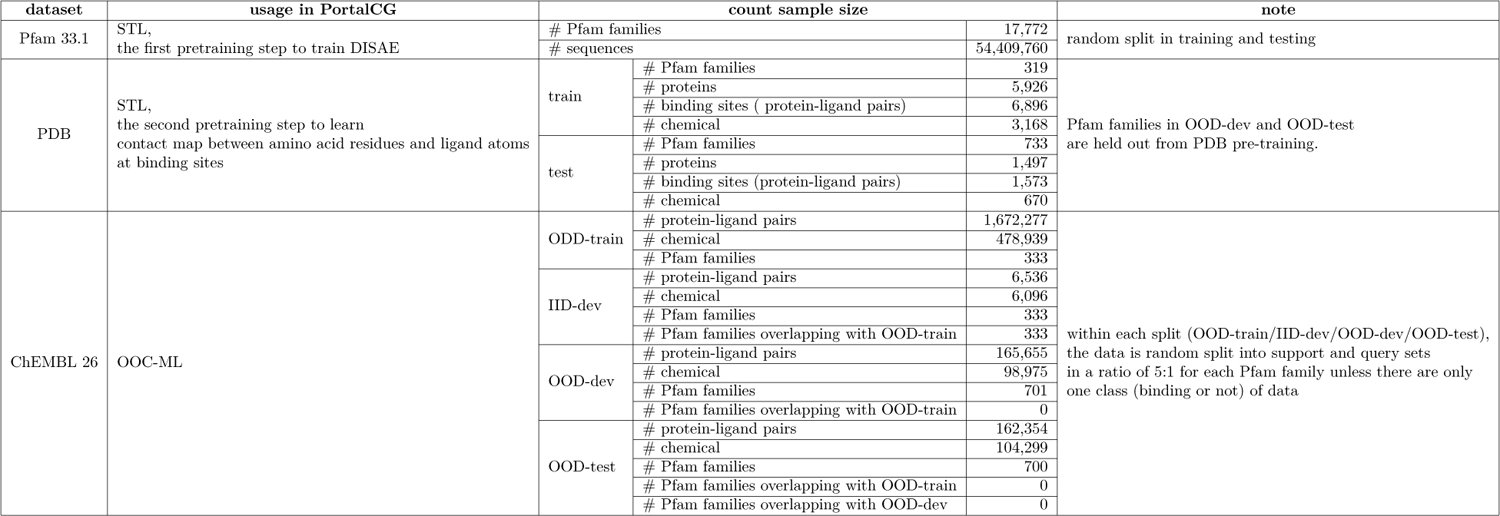
Data statistics for each training stage

65 compounds were synthesized for testing DRD1/2/3 binding activities. The procedures for the compound synthesis were detailed in Supplemental material Section 1.8, Scheme 1-5. DRD3 binding assays and Ki determinations were performed by the Psychoactive Drug Screening Program (PDSP).

For illuminating undruggable human proteins, around 6,000 drugs are collected from CLUE[39]. 12,475 undruggable proteins are collected by removing the druggable proteins in Pharos [37] and Casas’s druggable proteins [38] from human disease associated genes [17].

### 4.2 Algorithm

#### 4.2.1 Chemical representation

A chemical was represented as a graph and its embedding was learned using Graph Isomorphism Network (GIN)[50] which was designed to maximize the representational (or discriminative) power of a Graph Neural Network (GNN) based on Weisfeiler-Lehman (WL) graph isomorphism test. GIN is a common choice as chemical descriptor[35].

#### 4.2.2 Protein sequence pre-training

Protein descriptor is pretrained from scratch following exactly DISAE [1] on whole Pfam families, making it a universal protein language model. DISAE was inspired by recent success in self-supervised learning of unlabeled data in Nature Language Processing (NLP). It features a novel method, DIstilled Sequence Alignment Embedding (DISAE), for the protein sequence representation. DISAE can utilize all protein sequences to capture functional information without the knowledge of their structure and function. By incorporating biological knowledge into the sequence representation, DISAE can learn functionally important information about protein families that span a wide range of protein space. Different from existing sequence pre-training strategy that uses original protein sequences as input [22], DISAE distilled the original sequence into an ordered list of triplets by extracting evolutionary important positions from a multiple sequence alignment including insertions and deletions. Then long-range residue-residue interactions can be learned via the Transformer module in ALBERT[10]. A self-supervised masked language modeling (MLM) approach was used at this stage. In the MLM, 15% triplets are randomly masked and assumed that they are unknown. Then the remaining triplets are used to predict what the masked triplets are.

#### 4.2.3 Protein structure regularization

With the protein descriptor pretrained using the sequences from the whole Pfam, chemical descriptors and a distance learner were plugged in to fine-tune the protein representation. The distance learner follows Alphafold[4] which formulates a multi-way classification on a distrogram. Based on the histogram of distances between amino acids and ligand atoms, a histogram equalization^1^ was applied to formulate a 10-way classification on our binding site structure data as in Supplemental material Figure S9. Since protein and chemical descriptors output position-specific embeddings of a distilled protein sequence and all atoms of a chemical, pair-wise interaction features on the binding sites were created with a simple vector operation: a matrix multiplication was used to select embedding vectors of each binding residue and atom; multiply and broadcast the selected embedding vectors into a symmetric tensor as shown in the following, where *H* is embedding matrix of size (*number_of_residues, embedding_d_imension*) or (*number_of_atoms, embedding_d_imension*) and *A* is selector matrix[51],

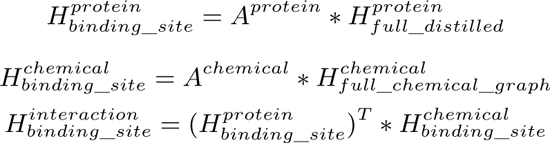

This pair-wise interaction feature tensor 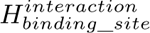 was fed into a Attentive Pooling[52] layer followed by feed-forward layer for final 10-way classification. Detailed model architecture configuration could be found in Supplemental Table S1 and Figure6. The intuition for the simplest form of distance learner is to put all stress of learning on the shared protein and chemical descriptors which will carry information across the end-to-end neural network. Again, with standard Adam optimization, shifted evaluation was used to select the “best” instance. Two versions of distance structure prediction were implemented, one formulated as a binary classification, i.e. contact prediction, one formulated as a multi-way classification, i.e. distogram prediction. The performance of the two version are similar, as shown in Figure S9.

#### 4.2.4 Out-of-cluster Meta Learning (OOC-ML)

With fine-tuned protein descriptor in the protein function space, a binary classifier is plugged on, which is a ResNet[53] layered with two linear layers as shown in Supplemental Table S1 and Figure6. What plays the major role in this phase is the optimization algorithm OOC-ML as shown in pseudocode **Algorithm 1** and Figure 1. The first level (low leverl) model training is reflected in line 4-9, and line 10 shows ensemble training of the second level (high level) models. Note that more variants could be derived from changing sampling rule (line 3 and 5) and the second level ensemble rule.

##### Algorithm 1: PortaCG, Out-of-cluster Meta-learning

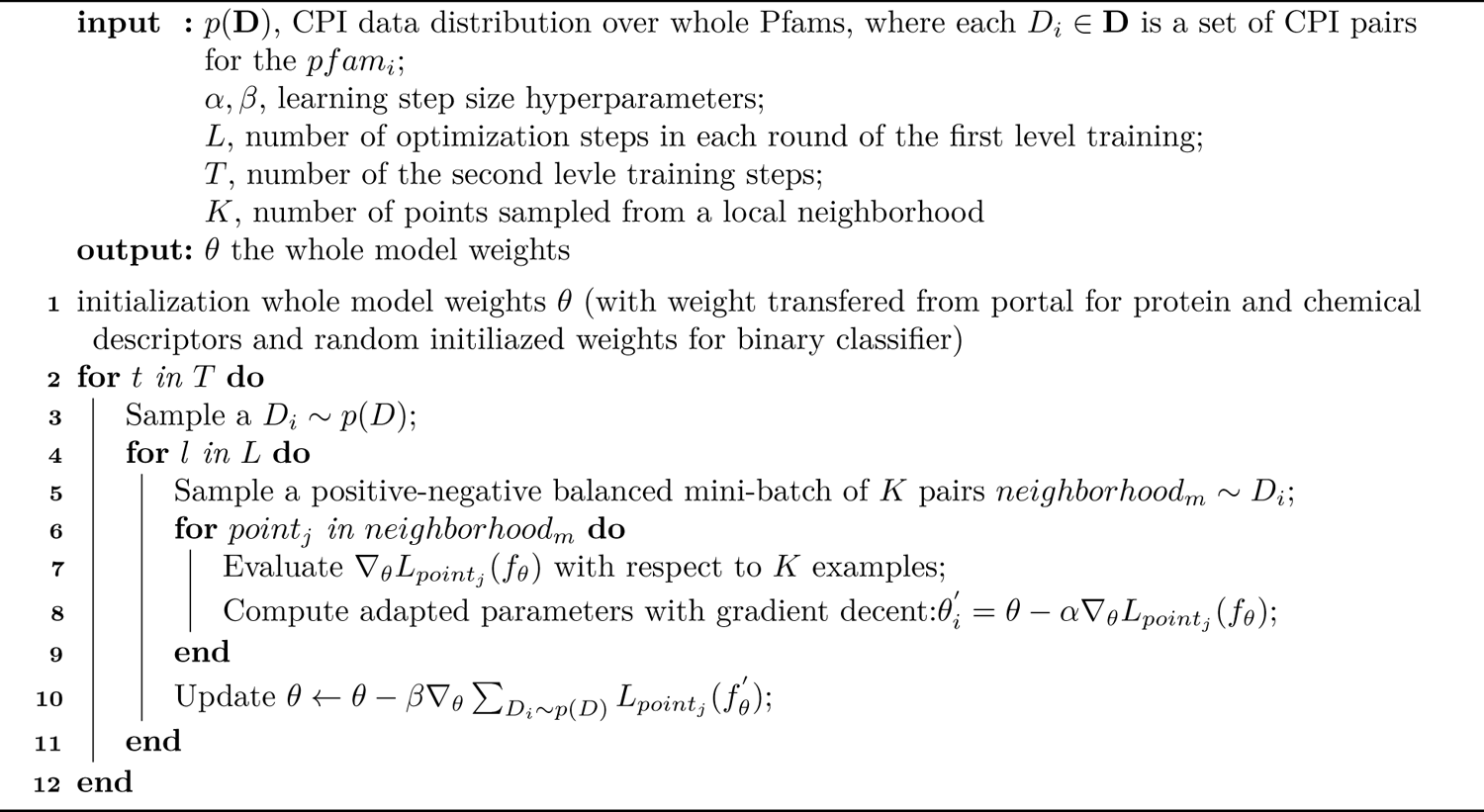

#### 4.2.5 Stress model instance selection

In classic training scheme common practice, there are 3-split data sets, “train set”, “dev set” and “test set”. Train set as the name suggested is used to train model. Test set as commonly expected is used to set an expectation of performance when applying the trained model to unseen data. Dev set is to select the preferred model instance. In OOD setting, data is split (see Table 1) such that dev set is a OOD from train set and test set is a OOD from both train and dev set. Deployment gap is calculated by deducting ODD-dev performance with OOD-test performance.

### 4.3 Baseline models

Machine learning methods for CPI predictions have been widely explored by many paradigms. As summarized in the survey [54], in addition to deep learning methods, there are similarity/distance based methods, matrix factorization, network-based, and feature-based methods. For CPI predictions with OOD generalization challenge, the similarity/distance-based, matrix factorization, and network-based methods have major obstacles. The Similarity/distance based methods rely on drug-drug similarity matrix and target-target similarity matrix as input. Because the similarities between dark proteins and proteins with known ligands are low, no reliable predictions can be made. Matrix Factorization is popular for its high efficiency but the cold-start nature of the dark proteins doesn’t fit matrix factorization paradigm. Network-based methods usually utilize protein-protein interactions. Such methods have advantages to predict the functional associations of ligand binding, but not the direct physical interactions. Furthermore, these methods are not scalable to millions of proteins and millions of chemicals. Almost of all studies based on these methods only focus on thousands of targets and thousands of drugs. PortalCG belongs to the category of the feature-based method. In our recently published work[1], we have shown that DISAE outperforms other state-of-the-art feature-based methods. Thus, we only compare PortalCG with DISAE in this paper.

Besides the machine learning method, protein-ligand docking (PLD) is a widely used approach to predict CPIs. We evaluate the performance of PLD based on Autodock Vina[28] and AlphaFold2 predicted structures[5] followed by SIGN re-scoring[29]. Structure-aware Interactive Graph Neural Networks (SIGN) [29] is a graph neural network proposed for the prediction of protein-ligand binding affinity. SIGN builds directional graphs to model the structures and interactions in protein-ligand complexes. Both distances and angles are integrated in the aggregation processes. SIGN is trained on PDBbind [55], which is a well-known public dataset containing 3D structures of protein-ligand complexes together with experimentally determined binding affinities. Similar to SIGN [29], we used the PDBbind v2016 dataset and the corresponding refined set, which contains 3767 complexes, to perform the training. We followed SIGN [29] for training and testing. For the directional graph used in SIGN, we constructed them with cutoff-threshold *θ_d_* = 5Å. The number of hidden layers is set to 2. All of the other settings are kept the same as those used in the original paper of SIGN. We randomly split the PDBbind refined set with a ratio of 9:1 for training and validation.

## Author Contributions

TC conceived the concept of PortalCG, implemented the algorithms, performed the experiments, and wrote the manuscript; Li Xie and SZ prepared data, performed the experiments, and wrote the manuscript; MC, DH, and YL implemented algorithms; KN, MD, and WWH designed and synthesized compounds; CM and PEB refined the concepts and wrote the manuscript; Lei Xie conceived and planned the experiments, wrote the manuscript.

## Data and software availability

Data used are described in section 4.3 and can be downloaded from public resource. Trained PortalCG model and PortalCG codes can be found in the Code Ocean. https://github.com/XieResearchGroup/PortalLearning.

## Acknowledgement

This project has been funded with federal funds from the National Institute of General Medical Sciences of National Institute of Health (R01GM122845) and the National Institute on Aging of the National Institute of Health (R01AD057555). We appreciate that Hansaim Lim helped with proof reading and provided constructive suggestions. Ki determinations, and receptor binding and activity profiles were generously provided by the National Institute of Mental Health’s Psychoactive Drug Screening Program, Contract #HHSN-271-2008-00025-C (NIMH PDSP). The NIMH PDSP is directed by Bryan L. Roth MD, PhD at the University of North Carolina at Chapel Hill and Project Officer Jamie Driscol at NIMH, Bethesda MD, USA.

## Footnotes

1 Histogram equalization: https://en.wikipedia.org/wiki/Histogram_equalization

## Notes

### Competing Interest Statement

The authors have declared no competing interest.

### Summary of Updates

Changed author list

## References

[1] T. Cai, H. Lim, K. A. Abbu, Y. Qiu, R. Nussinov, and L. Xie, “Msa-regularized protein sequence transformer toward predicting genome-wide chemical-protein interactions: Application to gpcrome deorphanization,” Journal of Chemical Information and Modeling, vol. 61, no. 4, pp. 1570–1582, 2021.

[2] J. Ma, S. H. Fong, Y. Luo, C. J. Bakkenist, J. P. Shen, S. Mourragui, L. F. Wessels, M. Hafner, R. Sharan, J. Peng, et al., “Few-shot learning creates predictive models of drug response that translate from high-throughput screens to individual patients,” Nature Cancer, vol. 2, no. 2, pp. 233–244, 2021.

[3] D. He, Q. Liu, Y. Wu, and L. Xie, “A context-aware deconfounding autoencoder for robust prediction of personalized clinical drug response from cell-line compound screening,” Nature Machine Intelligence, pp. 1–14, 2022.

[4] N. Hiranuma, H. Park, M. Baek, I. Anishchenko, J. Dauparas, and D. Baker, “Improved protein structure refinement guided by deep learning based accuracy estimation,” Nature communications, vol. 12, no. 1, pp. 1–11, 2021.

[5] J. Jumper, R. Evans, A. Pritzel, T. Green, M. Figurnov, O. Ronneberger, K. Tunyasuvunakool, R. Bates, A. Žídek, A. Potapenko, et al., “Highly accurate protein structure prediction with alphafold,” Nature, pp. 1–11, 2021.

[6] M. Baek, F. DiMaio, I. Anishchenko, J. Dauparas, S. Ovchinnikov, G. R. Lee, J. Wang, Q. Cong, L. N. Kinch, R. D. Schaeffer, et al., “Accurate prediction of protein structures and interactions using a 3-track network,” bioRxiv, 2021.

[7] Y. Li, P. Luo, Y. Lu, and F.-X. Wu, “Identifying cell types from single-cell data based on similarities and dissimilarities between cells,” BMC bioinformatics, vol. 22, no. 3, pp. 1–18, 2021.

[8] B. Schölkopf, F. Locatello, S. Bauer, N. R. Ke, N. Kalchbrenner, A. Goyal, and Y. Bengio, “Toward causal representation learning,” Proceedings of the IEEE, vol. 109, no. 5, pp. 612–634, 2021.

[9] W. Chen, Z. Yu, Z. Wang, and A. Anandkumar, “Automated synthetic-to-real generalization,” in International Conference on Machine Learning, pp. 1746–1756, PMLR, 2020.

[10] Z. Lan, M. Chen, S. Goodman, K. Gimpel, P. Sharma, and R. Soricut, “Albert: A lite bert for self-supervised learning of language representations,” arXiv preprint arXiv:1909.11942, 2019.

[11] C. Finn, P. Abbeel, and S. Levine, “Model-agnostic meta-learning for fast adaptation of deep networks,” CoRR,vol. abs/1703.03400, 2017.

[12] T. M. Hospedales, A. Antoniou, P. Micaelli, and A. J. Storkey, “Meta-learning in neural networks: A survey,” CoRR, vol. abs/2004.05439, 2020.

[13] T. I. Oprea, “Exploring the dark genome: implications for precision medicine,” Mammalian Genome, vol. 30, no. 7, pp. 192–200, 2019.

[14] G. Kustatscher, T. Collins, A.-C. Gingras, T. Guo, H. Hermjakob, T. Ideker, K. S. Lilley, E. Lundberg, E. M. Marcotte, M. Ralser, et al., “Understudied proteins: opportunities and challenges for functional proteomics,” Nature Methods, pp. 1–6, 2022.

[15] G. Kustatscher, T. Collins, A.-C. Gingras, T. Guo, H. Hermjakob, T. Ideker, K. S. Lilley, E. Lundberg, E. M. Marcotte, M. Ralser, et al., “An open invitation to the understudied proteins initiative,” Nature Biotechnology,pp. 1–3, 2022.

[16] L. Xie, L. Xie, S. L. Kinnings, and P. E. Bourne, “Novel computational approaches to polypharmacology as a means to define responses to individual drugs,” Annual review of pharmacology and toxicology, vol. 52, pp. 361–379, 2012.

[17] J. Piñero, J. M. Ramírez-Anguita, J. Saüch-Pitarch, F. Ronzano, E. Centeno, F. Sanz, and L. I. Furlong, “The disgenet knowledge platform for disease genomics: 2019 update,” Nucleic Acids Research, vol. 48, p. D845–D855, 1 2020.

[18] M. Karimi, D. Wu, Z. Wang, and Y. Shen, “Deepaffinity: interpretable deep learning of compound–protein affinity through unified recurrent and convolutional neural networks,” Bioinformatics, vol. 35, no. 18, pp. 3329–3338, 2019.

[19] H. Öztürk, A. Özgür, and E. Ozkirimli, “Deepdta: deep drug–target binding affinity prediction,” Bioinformatics,vol. 34, no. 17, pp. i821–i829, 2018.

[20] J. Devlin, M.-W. Chang, K. Lee, and K. Toutanova, “Bert: Pre-training of deep bidirectional transformers for language understanding,” arXiv preprint arXiv:1810.04805, 2018.

[21] S. Sledzieski, R. Singh, L. Cowen, and B. Berger, “Sequence-based prediction of protein-protein interactions: a structure-aware interpretable deep learning model,” bioRxiv, 2021.

[22] A. Rives, J. Meier, T. Sercu, S. Goyal, Z. Lin, J. Liu, D. Guo, M. Ott, C. L. Zitnick, J. Ma, et al., “Biological structure and function emerge from scaling unsupervised learning to 250 million protein sequences,” Proceedings of the National Academy of Sciences, vol. 118, no. 15, p. e2016239118, 2021.

[23] L. Xie and P. E. Bourne, “Detecting evolutionary relationships across existing fold space, using sequence order-independent profile–profile alignments,” Proceedings of the National Academy of sciences, vol. 105, no. 14, pp. 5441–5446, 2008.

[24] M. AlQuraishi and P. K. Sorger, “Differentiable biology: using deep learning for biophysics-based and data-driven modeling of molecular mechanisms,” Nature methods, vol. 18, no. 10, pp. 1169–1180, 2021.

[25] J. Mistry, S. Chuguransky, L. Williams, M. Qureshi, G. A. Salazar, E. L. Sonnhammer, S. C. Tosatto, L. Paladin, S. Raj, L. J. Richardson, et al., “Pfam: The protein families database in 2021,” Nucleic Acids Research, vol. 49, no. D1, pp. D412–D419, 2021.

[26] A. Gaulton, A. Hersey, M. Nowotka, A. P. Bento, J. Chambers, D. Mendez, P. Mutowo, F. Atkinson, L. J. Bellis, E. Cibrián-Uhalte, M. Davies, N. Dedman, A. Karlsson, M. P. Magariños, J. P. Overington, G. Papadatos, I. Smit, and A. R. Leach, “The ChEMBL database in 2017,” Nucleic Acids Research, vol. 45, pp. D945–D954, 11 2016.

[27] H. Huang, G. Zhang, Y. Zhou, C. Lin, S. Chen, Y. Lin, S. Mai, and Z. Huang, “Reverse screening methods to search for the protein targets of chemopreventive compounds,” Frontiers in chemistry, vol. 6, p. 138, 2018.

[28] O. Trott and A. J. Olson, “Autodock vina: improving the speed and accuracy of docking with a new scoring function, efficient optimization and multithreading,” Journal of Computational Chemistry, vol. 31, pp. 455–461, 2010.

[29] S. Li, J. Zhou, T. Xu, L. Huang, F. Wang, H. Xiong, W. Huang, D. Dou, and H. Xiong, “Structure-aware interactive graph neural networks for the prediction of protein-ligand binding affinity,” in Proceedings of the 27th ACM SIGKDD Conference on Knowledge Discovery & Data Mining, pp. 975–985, 2021.

[30] S. Z. Grinter and X. Zou, “Challenges, applications, and recent advances of protein-ligand docking in structure-based drug design,” Molecules, vol. 19, no. 7, pp. 10150–10176, 2014.

[31] M. Jaiteh, I. Rodríguez-Espigares, J. Selent, and J. Carlsson, “Performance of virtual screening against gpcr homology models: Impact of template selection and treatment of binding site plasticity,” PLoS computational biology, vol. 16, no. 3, p. e1007680, 2020.

[32] M. M. Mysinger, M. Carchia, J. J. Irwin, and B. K. Shoichet, “Directory of useful decoys, enhanced (dud-e): better ligands and decoys for better benchmarking,” Journal of medicinal chemistry, vol. 55, no. 14, pp. 6582–6594, 2012.

[33] S. T. Ewing, C. Dorcely, R. Maidi, G. Paker, E. Schelbaum, and R. Ranaldi, “Low-dose polypharmacology targeting dopamine d1 and d3 receptors reduces cue-induced relapse to heroin seeking in rats,” Addiction Biology, vol. 26, no. 4, p. e12988, 2021.

[34] E. Galaj, S. Ewing, and R. Ranaldi, “Dopamine d1 and d3 receptor polypharmacology as a potential treatment approach for substance use disorder,” Neuroscience & Biobehavioral Reviews, vol. 89, pp. 13–28, 2018.

[35] W. Hu, B. Liu, J. Gomes, M. Zitnik, P. Liang, V. Pande, and J. Leskovec, “Strategies for pre-training graph neural networks” 2020.

[36] C. Finan, A. Gaulton, F. A. Kruger, R. T. Lumbers, T. Shah, J. Engmann, L. Galver, R. Kelley, A. Karlsson, R. Santos, et al., “The druggable genome and support for target identification and validation in drug development,” Science translational medicine, vol. 9, no. 383, 2017.

[37] T. K. Sheils, S. L. Mathias, K. J. Kelleher, V. B. Siramshetty, D.-T. Nguyen, C. G. Bologa, L. J. Jensen, D. Vidović, A. Koleti, S. C. Schürer, A. Waller, J. J. Yang, J. Holmes, G. Bocci, N. Southall, P. Dharkar, E. Mathé, A. Simeonov, and T. I. Oprea, “Utcrd and pharos 2021: mining the human proteome for disease biology,” Nucleic Acids Research, vol. 49, pp. D1334–D1346, 1 2021.

[38] C. Finan, A. Gaulton, F. A. Kruger, R. T. Lumbers, T. Shah, J. Engmann, L. Galver, R. Kelley, A. Karlsson, R. Santos, J. P. Overington, A. D. Hingorani, and J. P. Casas, “The druggable genome and support for target identification and validation in drug development,” Science Translational Medicine, vol. 9, p. eaag1166, 3 2017.

[39] S. M. Corsello, J. A. Bittker, Z. Liu, J. Gould, P. McCarren, J. E. Hirschman, S. E. Johnston, A. Vrcic, B. Wong, M. Khan, et al., “The drug repurposing hub: a next-generation drug library and information resource,” Nature medicine, vol. 23, no. 4, pp. 405–408, 2017.

[40] X. Jiao, B. T. Sherman, D. W. Huang, M. W. B. Robert Stephens, H. C. Lane, and R. A. Lempicki, “David-ws: a stateful web service to facilitate gene/protein list analysis,” Bioinformatics, vol. 28, p. 1805–1806, 7 2012.

[41] D. O. Bates, J. C. Morris, S. Oltean, and L. F. Donaldson, “Pharmacology of modulators of alternative splicing,” Pharmacological reviews, vol. 69, no. 1, pp. 63–79, 2017.

[42] K.-q. Le, B. S. Prabhakar, W.-j. Hong, and L.-c. Li, “Alternative splicing as a biomarker and potential target for drug discovery,” Acta Pharmacologica Sinica, vol. 36, no. 10, pp. 1212–1218, 2015.

[43] J. E. Love, E. J. Hayden, and T. T. Rohn, “Alternative splicing in alzheimer’s disease,” Journal of Parkinson’s disease and Alzheimer’s disease, vol. 2, no. 2, 2015.

[44] S. Zhang, Y. Liu, and L. Xie, “Efficient and accurate physics-aware multiplex graph neural networks for 3d small molecules and macromolecule complexes,” arXiv preprint arXiv:2206.02789, 2022.

[45] Y. Liu, H. Lim, and L. Xie, “Exploration of chemical space with partial labeled noisy student self-training and self-supervised graph embedding,” BMC bioinformatics, vol. 23, no. 3, pp. 1–21, 2022.

[46] Y. Liu, Y. Wu, X. Shen, and L. Xie, “Covid-19 multi-targeted drug repurposing using few-shot learning,” Frontiers in Bioinformatics, vol. 1, 2021.

[47] H. M. Berman, J. Westbrook, Z. Feng, G. Gilliland, T. N. Bhat, H. Weissig, I. N. Shindyalov, and P. E. Bourne, “The Protein Data Bank,” Nucleic Acids Research, vol. 28, pp. 235–242, 01 2000.

[48] J. Yang, A. Roy, and Y. Zhang, “Biolip: a semi-manually curated database for biologically relevant ligand–protein interactions,” Nucleic acids research, vol. 41, no. D1, pp. D1096–D1103, 2012.

[49] S. C. Potter, A. Luciani, S. R. Eddy, Y. Park, R. Lopez, and R. D. Finn, “Hmmer web server: 2018 update,” Nucleic acids research, vol. 46, no. W1, pp. W200–W204, 2018.

[50] K. Xu, W. Hu, J. Leskovec, and S. Jegelka, “How powerful are graph neural networks?,” arXiv preprint arXiv:1810.00826, 2018.

[51] S. Boyd and L. Vandenberghe, Introduction to applied linear algebra: vectors, matrices, and least squares. Cambridge university press, 2018.

[52] C. d. Santos, M. Tan, B. Xiang, and B. Zhou, “Attentive pooling networks,” arXiv preprint arXiv:1602.03609, 2016.

[53] K. He, X. Zhang, S. Ren, and J. Sun, “Deep residual learning for image recognition,” in Proceedings of the IEEE conference on computer vision and pattern recognition, pp. 770–778, 2016.

[54] M. Bagherian, E. Sabeti, K. Wang, M. A. Sartor, Z. Nikolovska-Coleska, and K. Najarian, “Machine learning approaches and databases for prediction of drug–target interaction: a survey paper,” Briefings in bioinformatics,vol. 22, no. 1, pp. 247–269, 2021.

[55] R. Wang, X. Fang, Y. Lu, and S. Wang, “The pdbbind database: Collection of binding affinities for protein-ligand complexes with known three-dimensional structures,” Journal of medicinal chemistry, vol. 47, no. 12, pp. 2977–2980, 2004.

