## Supplemental for "Binding Site-enhanced Sequence Pretraining and Out-of-cluster Meta-learning Predict Genome-Wide Chemical-Protein Interactions for Dark Proteins"

**Reference** **30**

### 1 Methods

In this section, we present the detailed methodology of PortalCG.

#### 1.1 Implementation details

Four major variants of models are trained as shown in main content Table 3 for controlled factor experiments to verify the contribution of key components of PortalCG. In this section we present implementation details.

Due to the large number of total samples, all training are carried out under global step-based formalization instead of epoch-based. Typically, a deep learning model is trained for numerous epochs, in each epoch the model will loop over all training data. Evaluation will be carried out once on the whole test data set at the end of each epoch. In the global step formalization, a mini-batch is sampled at uniform random from pre-split training data set. For a pre-defined total number of global steps, this mini-batch sampling will be repeated. To evaluate along the way of training, for every $m$ global steps of training, a subset of OOD-test data is sampled uniformly randomly from a pre-split test set. To compute generalization gaps, in addition to evaluate on OOD-test set split according to the shifted evaluation, an OOD-dev set and and IID-dev are held out comparing to the train set distribution for the evaluation as well.

#### 1.2 Evaluation metrics

Distogram prediction uses an average accuracy on the distogram. CPI binary classification uses F1, ROC-AUC and PR-AUC for overall evaluation with breakdown by class F1, recall and precision scores.

#### 1.3 Docking as baseline

Protein-ligand docking was performed using Autodock Vina[1]. In the benchmark study, the whole protein surface search implemented in the Autodock Vina was applied to identify the ligand binding pocket. The center of each protein was set as the center of the binding pocket. For the DRD1/DRD2/DRD3, the center of the co-crystallized ligand was set as the center of the binding pocket. The largest distance of the protein atoms to the center of the protein is calculated for each x, y, and z direction to define the edge of each protein. 10 Angstrom of extra space was added to the protein edge to set up the search space for the docking. We selected the 9 binding poses that had the best scores.

#### 1.4 Production level for deployment

To create a production level model, three models were trained in PortalCG with only difference in data split. Dev set was OOD in respect of training set to make sure there was no overlapped Pfam families between them. By rotating Pfam families between training set and OOD-dev set in the fashion of a cross-validation, each of the three models was trained on different train set in light of Pfam families involved. Then a voting mechanism was used to make the final prediction.

#### 1.5 Additional results in dark chemical genomics space exploration

When we consider the proteins in Tbio, there are 9545 proteins which are not int Casas's druggable proteins. If 0.67 was used as the cutoff, 219 proteins were predicted as positive hits. The gene enrichment analysis result for these proteins was listed in Table S7. Disease associated with these

219 human proteins were also listed in main context Table 5. Since one protein is always related with multiple diseases, these diseases are ranked by the number of their associated proteins and the top 10 diseases were listed in the table. Most of top ranked diseases are related with cancer development. 21 drugs that are approved or in clinical trial are predicted to interact with these proteins as shown in Table S8.

If the proteins in Tbio were removed from the undruggable list, only 2930 proteins were left. If 0.67 was used as the cutoff, there will be only 41 proteins predicted positive and no significant enrichment with David gene enrichment analysis. So 0.665 was used as a cutoff, and 348 proteins were predicted as positive hits. The gene enrichment analysis result for these proteins was listed in Table S9. Disease associated with these 348 human proteins were also listed in Table S10. 42 drugs that are approved or in clinical trial are predicted to interact with these proteins as shown in Table S11.

#### 1.6 Compound design and synthesis

5 series of compounds were designed and synthesized as detailed below.

##### 1.6.1 Serial 1 compounds

Serial 1 compounds were synthesized as shown in Scheme 1 and as described in reference [2]. Briefly, the brominated phenol tetrahydroprotoberberine derivative 1 was alkylated with various alkyl halides to install the C9 alkoxy groups in compounds 2a-2g. Debenzylation of compounds 2a-2g via treatment with ethanolic HCl afforded analogs 3a-3g. Removal of the bromine group in compounds 2a-2g gave compounds 4a-4g, which were subsequently debenzylated to afford analogs 5a-5g.

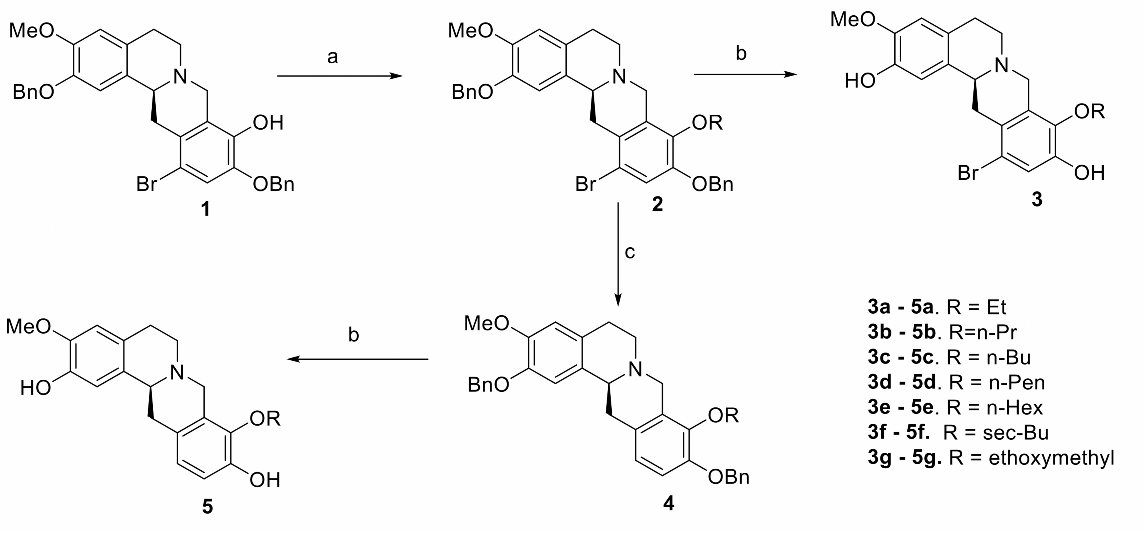

**Scheme 1**. *Reagents and condition:* a) appropriate alkyl halide, $K_2CO_3$, DMF, 2-4h; b) conc. HCl, EtOH, 70 °C, 1.5 – 2 h; c) i-PrMgCl, LiCl, THF, 0 °C, 2 h

##### 1.6.2 Serial 2 compounds

The synthesis of serial 2 compounds is shown in Scheme 2 and is described in detail in reference [?]. Briefly, the benzylated tetrahydroisoquinoline 6 was debenzylated to give compound 7. Treatment of the phenol 6 with appropriate alkyl halides allowed for alkylation of the C10 phenol group of 6. The intermediate C10 alkoxy derivatives were then debenzylated to afford the C10 alkoxylated/C2 phenol derivatives 8a – 8h.

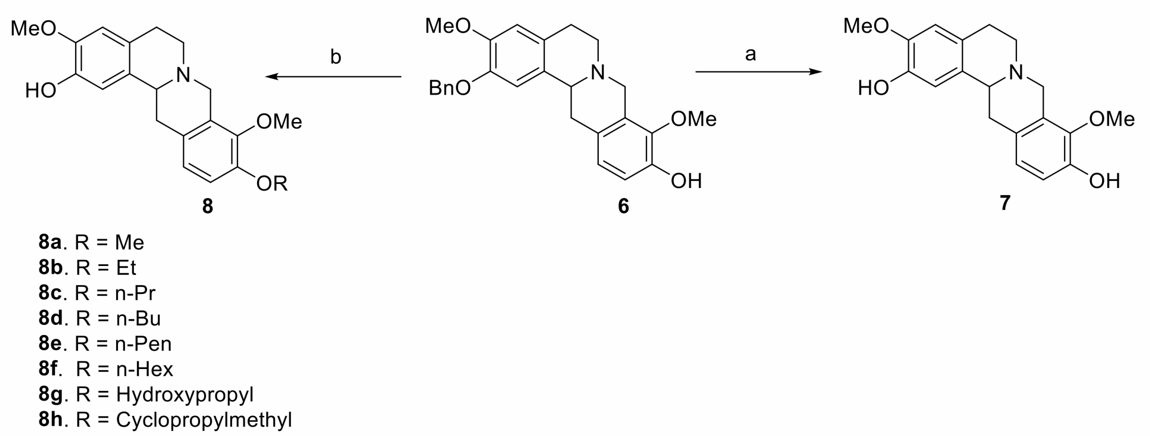

**Scheme 2**. *Reagents and conditions*: conc. HCl, MeOH, reflux; b) (i) appropriate akyl halide, DMF, $CH_3CN$, rt to reflux; (ii) conc. HCl, MeOH, reflux.

##### 1.6.3 Serial 3 compounds

Serial 3 ompounds were synthesized as depicted in Scheme 3. Further details on the synthesis of the analogs and precursors may be found in reference [3]. The acetate group in compound 9 was hydrolyzed to an intermediate alcohol, which was subsequently reacted with thionyl chloride. The alkyl chloride thus formed underwent ring closure to form tetrahydroprotoberberine 10. The C3 phenol of 10 was methylated and the C2 and C10 benzyl groups of the resulting compounds were removed under acidic conditions to afford analogs 11a-11e.

##### 1.6.4 Serial 4 compounds

Synthesis of the serial 4 compounds is shown in Scheme 4 and reported in a previous publication [4]. The amine 12 was coupled to various carboxylic acids and the resulting benzylated amides were debenzylated by refluxing in ethanolic HCl, yielding analogs 13a-13m. The phenol 13m was methylated to provide analog 13n.

##### 1.6.5 Serial 5 compounds

Serial 5 compounds were synthesized as depicted in Scheme 5 and as previously reported in reference [5]. Briefly, amine 14a was coupled with various aromatic carboxylic acids to give intermediate amides which were subsequently debenzylated to give the corresponding amide analogs 15a-15r.

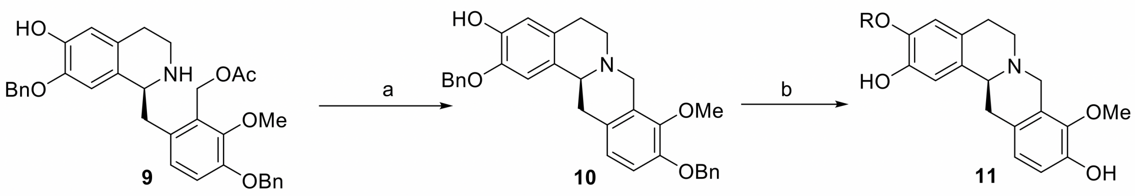

**11a**. R = Et
**11b**. R= n-Pr
**11c**. R = n-Bu
**11d**. R = n-Pen
**11e**. R = n-Hexyl

**Scheme 3**. *Reagents and conditions*: a) (i) 10% NaOH, EtOH, rt; (ii) $SOCl_2$, $CH_2Cl_2$, $NaHCO_3$; b) RBr, $K_2CO_3$, DMF, rt; (ii) conc. HCl, EtOH, reflux

Compounds 15s-15t were synthesized by amide coupling of amine 14b with the corresponding cyanophenyl carboxylic acids. Compounds 16a-16c were synthesized by treatment of corresponding precursors in the compound 15 series with BBr3, to effect demethylation to the catechol functionality.

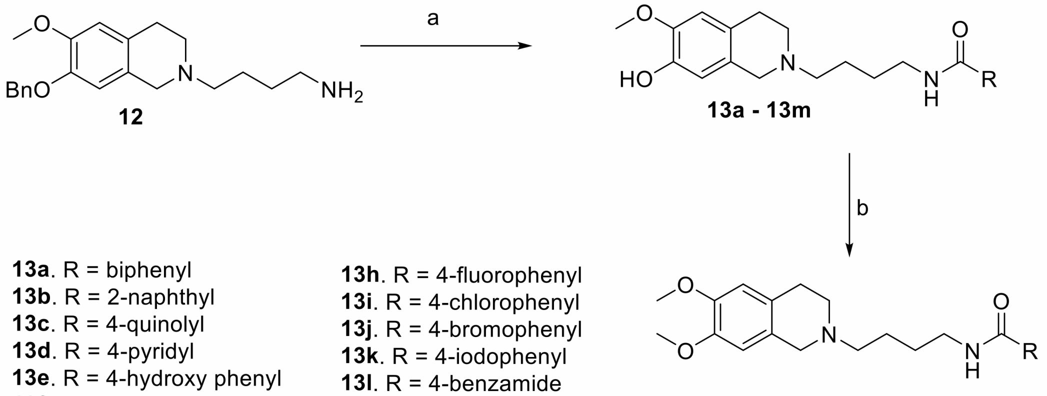

**13a**. R = biphenyl
**13b**. R = 2-naphthyl
**13c**. R = 4-quinolyl
**13d**. R = 4-pyridyl
**13e**. R = 4-hydroxy phenyl
**13f**. R = 4-methoxy phenyl
**13g**. R = 3-methoxyphenyl
**13h**. R = 4-fluorophenyl
**13i**. R = 4-chlorophenyl
**13j**. R = 4-bromophenyl
**13k**. R = 4-iodophenyl
**13l**. R = 4-benzamide
**13m**. R = 4-cyanobenzyl

**13n**. R = 4-cyanobenzyl

**Scheme 4**. *Reagents and conditions*: a) (i)RCOOH, EDC, DCM, rt; (ii) Conc. HCl, EtOH, reflux; b) **13m**, MeI, $K_2CO_3$, acetone

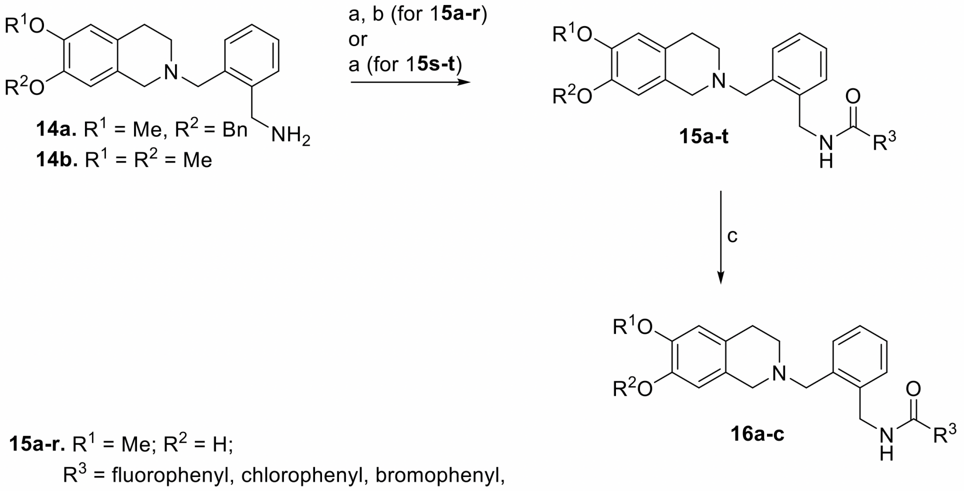

**14a.** $R^1$ = Me, $R^2$ = Bn
**14b.** $R^1$ = $R^2$ = Me

**15a-r.** $R^1$ = Me; $R^2$ = H;
$R^3$ = fluorophenyl, chlorophenyl, bromophenyl, iodophenyl, methoxyphenyl, cyanophenyl, 2-naphthyl, biphenyl
**15s.** $R^1$ = $R^2$ = Me; $R^3$ = 3-cyanophenyl
**15t.** $R^1$ = $R^2$ = Me; $R^3$ = 4-cyanophenyl

**16a**. $R^1$ = $R^2$ = H; $R^3$ = 3-bromophenyl
**16b**. $R^1$ = $R^2$ = H; $R^3$ = 3-cyanophenyl
**16c**. $R^1$ = $R^2$ = H; $R^3$ = 4-cyanophenyl

**Scheme 5**. *Reagents and conditions*: (a) HBTU, appropriate carboxylic acid, TEA, 12 h, rt; (b) conc. HCl, $CH_3COOH$, 12 h, 40 °C; (c) $BBr_3$, DCM, 0°C, 2h

### 2 Additional tables

Table S 1: Model architecture configuration

| | | |
|---|---|---|
| Protein descriptor | layers | Albert –>Resnet |
| | embedding dimension | 256 |
| Chemical descriptor | backbone | GIN |
| | number of layers | 5 |
| | embedding dimension | 300 |
| | aggregation methods | sum |
| | drop out ratio | 0.5 |
| Interaction learner | layers | Attentive pooling –>2 layers of MLP |
| | embedding dimension | 128 |
| Structure residue-atom pair wise feature learner | layers | matrix multiplication of protein and chemical embedding vectors |
| Classifier | layers | 2 layers of MLP |
| | embedding dimension | 64 |

Table S 2: 65 compounds tested for selective dual DRD1/3 antagonists. na: no detectable binding affinity.

| SMILES | D1R Ki in nM | D3R Ki in nM | D2R Ki in nM |
|---|---|---|---|
| BrC1=CC(O)=C(OCC)C2=C1C[C@H]3C4=CC(O)=C(OC)C=C4CCN3C2 | 1.3 | 373 | na |
| BrC1=CC(O)=C(OCCC)C2=C1C[C@H]3C4=CC(O)=C(OC)C=C4CCN3C2 | 2.2 | 195 | 70 |
| BrC1=CC(O)=C(OCCCC)C2=C1C[C@H]3C4=CC(O)=C(OC)C=C4CCN3C2 | 2.3 | na | 107 |
| BrC1=CC(O)=C(OCCCCC)C2=C1C[C@H]3C4=CC(O)=C(OC)C=C4CCN3C2 | 32 | 3605 | 398 |
| BrC1=CC(O)=C(OCCCCCC)C2=C1C[C@H]3C4=CC(O)=C(OC)C=C4CCN3C2 | 68 | 3171 | 731 |
| BrC1=CC(O)=C(OCC(C)C)C2=C1C[C@H]3C4=CC(O)=C(OC)C=C4CCN3C2 | 9.7 | 2070 | 331 |
| BrC1=CC(O)=C(OCCOC)C2=C1C[C@H]3C4=CC(O)=C(OC)C=C4CCN3C2 | 4.2 | 726 | na |
| OC1=C(OC)C=C2C([C@H](CC(C=CC(O)=C3OCC)=C3C4)N4CC2)=C1 | 5.3 | 106 | na |
| OC1=C(OC)C=C2C([C@H](CC(C=CC(O)=C3OCCC)=C3C4)N4CC2)=C1 | 12 | 175 | 41 |
| OC1=C(OC)C=C2C([C@H](CC(C=CC(O)=C3OCCCC)=C3C4)N4CC2)=C1 | 11 | 131 | na |
| OC1=C(OC)C=C2C([C@H](CC(C=CC(O)=C3OCCCCC)=C3C4)N4CC2)=C1 | 31 | 3213 | 23 |
| OC1=C(OC)C=C2C([C@H](CC(C=CC(O)=C3OCCCCCC)=C3C4)N4CC2)=C1 | 77 | 195 | na |
| OC1=C(OC)C=C2C([C@H](CC(C=CC(O)=C3OCC(C)C)=C3C4)N4CC2)=C1 | 65 | 450 | 104 |
| OC1=C(OC)C=C2C([C@H](CC(C=CC(O)=C3OCCOC)=C3C4)N4CC2)=C1 | 7.7 | 31 | 18 |
| OC1=C(OC)C=C2C([C@H](CC(C=CC(O)=C3OC)=C3C4)N4CC2)=C1 | 5.9 | 30 | 974 |
| OC1=C(OC)C=C2C([C@H](CC(C=CC(OC)=C3OC)=C3C4)N4CC2)=C1 | 15 | 37 | 102 |
| OC1=C(OC)C=C2C([C@H](CC(C=CC(OCC)=C3OC)=C3C4)N4CC2)=C1 | 124 | 44 | 159 |
| OC1=C(OC)C=C2C([C@H](CC(C=CC(OCCC)=C3OC)=C3C4)N4CC2)=C1 | 7.8 | 53 | 99 |
| OC1=C(OC)C=C2C([C@H](CC(C=CC(OCCCC)=C3OC)=C3C4)N4CC2)=C1 | 11 | 65 | 121 |
| OC1=C(OC)C=C2C([C@H](CC(C=CC(OCCCCC)=C3OC)=C3C4)N4CC2)=C1 | 19 | 101 | 283 |
| OC1=C(OC)C=C2C([C@H](CC(C=CC(OCCCCCC)=C3OC)=C3C4)N4CC2)=C1 | 40 | 220 | na |
| OC1=C(OC)C=C2C([C@H](CC(C=CC(OCCCO)=C3OC)=C3C4)N4CC2)=C1 | 6.7 | 110 | 211 |
| OC1=C(OC)C=C2C([C@H](CC(C=CC(OCC3CC3)=C4OC)=C4C5)N5CC2)=C1 | 12 | 54 | 54 |
| OC1=C(OCC)C=C2C([C@H](CC(C=CC(O)=C3OC)=C3C4)N4CC2)=C1 | 17 | 40 | 2089 |
| OC1=C(OCCC)C=C2C([C@H](CC(C=CC(O)=C3OC)=C3C4)N4CC2)=C1 | 15 | 46 | 2682 |
| OC1=C(OCCCC)C=C2C([C@H](CC(C=CC(O)=C3OC)=C3C4)N4CC2)=C1 | 53 | 51 | 2739 |
| OC1=C(OCCCCC)C=C2C([C@H](CC(C=CC(O)=C3OC)=C3C4)N4CC2)=C1 | 31 | 33 | 1433 |
| OC1=C(OCCCCCC)C=C2C([C@H](CC(C=CC(O)=C3OC)=C3C4)N4CC2)=C1 | 42 | 26 | 1491 |
| OC1=C(C=C2CCN(CC2=C1)CCCCNC(C3=CC=C(C4=CC=CC=C4)C=C3)=O)OC | 800 | 2.7 | 800 |
| OC1=C(C=C2CCN(CC2=C1)CCCCNC(C3=CC=C(C=CC=C4)C4=C3)=O)OC | 800 | 2.7 | 700 |
| OC1=C(C=C2CCN(CC2=C1)CCCCNC(C3=CC=NC4=C3C=CC=C4)=O)OC | 1810 | 22 | na |
| OC1=C(C=C2CCN(CC2=C1)CCCCNC(C3=CC=NC=C3)=O)OC | na | 24 | na |
| OC1=C(C=C2CCN(CC2=C1)CCCCNC(C3=CC=C(O)C=C3)=O)OC | na | 8.7 | na |
| OC1=C(C=C2CCN(CC2=C1)CCCCNC(C3=CC=C(OC)C=C3)=O)OC | na | 5.9 | na |
| OC1=C(C=C2CCN(CC2=C1)CCCCNC(C3=CC=C(F)C=C3)=O)OC | na | 4.4 | na |
| OC1=C(C=C2CCN(CC2=C1)CCCCNC(C3=CC=C(Cl)C=C3)=O)OC | 680 | 2.1 | 1100 |
| OC1=C(C=C2CCN(CC2=C1)CCCCNC(C3=CC=C(Br)C=C3)=O)OC | 500 | 3.4 | na |
| OC1=C(C=C2CCN(CC2=C1)CCCCNC(C3=CC=C(I)C=C3)=O)OC | 1600 | 10 | na |
| OC1=C(C=C2CCN(CC2=C1)CCCCNC(C3=CC=C(C(N)=O)C=C3)=O)OC | na | 28 | 860 |
| OC1=C(C=C2CCN(CC2=C1)CCCCNC(C3=CC=C(C#N)C=C3)=O)OC | na | 6.3 | na |
| OC1=C(C=C2CCN(CC2=C1)CCCCNC(C3=CC=CC(OC)=C3)=O)OC | na | 12 | na |
| O=C(C1=CC=C(C#N)C=C1)NCCCCN(CC2=C3)CCC2=CC(OC)=C3OC | na | 410 | na |
| OC1=C(C=C2CCN(CC2=C1)CC3=C(CNC(C4=CC=C(C5=CC=CC=C5)C=C4)=O)C=CC=C3)OC | na | na | na |
| OC1=C(C=C2CCN(CC2=C1)CC3=C(CNC(C4=CC=C(C=CC=C5)C5=C4)=O)C=CC=C3)OC | na | 780 | 310 |
| OC1=C(C=C2CCN(CC2=C1)CC3=C(CNC(C4=CC=CC=C4Cl)=O)C=CC=C3)OC | na | 380 | 190 |
| OC1=C(C=C2CCN(CC2=C1)CC3=C(CNC(C4=CC=CC=C4Br)=O)C=CC=C3)OC | na | 540 | 420 |
| OC1=C(C=C2CCN(CC2=C1)CC3=C(CNC(C4=CC=CC=C4OC)=O)C=CC=C3)OC | na | 120 | 78 |
| OC1=C(C=C2CCN(CC2=C1)CC3=C(CNC(C4=CC=CC(Cl)=C4)=O)C=CC=C3)OC | na | 140 | 100 |
| OC1=C(C=C2CCN(CC2=C1)CC3=C(CNC(C4=CC=CC(Br)=C4)=O)C=CC=C3)OC | na | 170 | 150 |
| OC1=C(C=C2CCN(CC2=C1)CC3=C(CNC(C4=CC=CC(OC)=C4)=O)C=CC=C3)OC | na | 280 | 140 |
| OC1=C(C=C2CCN(CC2=C1)CC3=C(CNC(C4=CC=CC(C#N)=C4)=O)C=CC=C3)OC | na | 26 | 39 |
| OC1=C(C=C2CCN(CC2=C1)CC3=C(CNC(C4=CC=C(F)C=C4)=O)C=CC=C3)OC | na | 140 | 52 |
| OC1=C(C=C2CCN(CC2=C1)CC3=C(CNC(C4=CC=C(Cl)C=C4)=O)C=CC=C3)OC | na | 55 | 47 |
| OC1=C(C=C2CCN(CC2=C1)CC3=C(CNC(C4=CC=C(Br)C=C4)=O)C=CC=C3)OC | na | 160 | 76 |
| OC1=C(C=C2CCN(CC2=C1)CC3=C(CNC(C4=CC=C(I)C=C4)=O)C=CC=C3)OC | na | na | 84 |
| OC1=C(C=C2CCN(CC2=C1)CC3=C(CNC(C4=CC=C(OC)C=C4)=O)C=CC=C3)OC | na | 180 | 74 |
| OC1=C(C=C2CCN(CC2=C1)CC3=C(CNC(C4=CC=C(C#N)C=C4)=O)C=CC=C3)OC | na | 310 | 190 |
| OC1=C(C=C2CCN(CC2=C1)CC3=C(CNC(C4=CC=CC(Cl)=C4Cl)=O)C=CC=C3)OC | na | 460 | 280 |
| OC1=C(C=C2CCN(CC2=C1)CC3=C(CNC(C4=CC=CC(OC)=C4OC)=O)C=CC=C3)OC | na | 57 | na |
| OC1=C(C=C2CCN(CC2=C1)CC3=C(CNC(C4=CC=C(I)C(OC)=C4)=O)C=CC=C3)OC | 1970 | 24 | 27 |
| O=C(C1=CC=CC(C#N)=C1)NCC(C=CC=C2)=C2CN(CC3=C4)CCC3=CC(OC)=C4OC | 510 | 1.2 | 46 |
| O=C(C1=CC=C(C#N)C=C1)NCC(C=CC=C2)=C2CN(CC3=C4)CCC3=CC(OC)=C4OC | 58 | 3.4 | 50 |
| OC(C=C1CCN(CC1=C2)CC3=C(CNC(C4=CC=C(C#N)C=C4)=O)C=CC=C3)=C2O | na | 28 | 83 |
| OC(C=C1CCN(CC1=C2)CC3=C(CNC(C4=CC=CC(C#N)=C4)=O)C=CC=C3)=C2O | na | 370 | 900 |
| OC(C=C1CCN(CC1=C2)CC3=C(CNC(C4=CC=CC(Br)=C4)=O)C=CC=C3)=C2O | 91 | 2 | 56 |

Table S 3: Undruggable human disease-associated proteins selected by ProtalCG.

| Uniprot | Protein name | Drug name |
|---|---|---|
| Q8TEX9 | Importin-4 | fenebrutinib |
| Q8NB66 | Protein unc-13 homolog C | AI-10-49 |
| Q9NZF1 | Placenta-specific gene 8 protein | AI-10-49 |
| Q96G03 | Phosphoglucomutase-2 | fenebrutinib |
| Q8IWU4 | Zinc transporter 8 | fenebrutinib |
| P53004 | Biliverdin reductase A | AI-10-49 |
| P40879 | Chloride anion exchanger | fenebrutinib |
| Q8IYL2 | Probable tRNA | fenebrutinib |
| Q9ULL4 | Plexin-B3 | AI-10-49 |
| Q96SZ6 | Mitochondrial tRNA methylthiotransferase CDK5RAP1 | fenebrutinib |
| Q9BRT9 | DNA replication complex GINS protein SLD5 | fenebrutinib |
| Q8N2U0 | Transmembrane protein 256 | CCT137690 |
| Q9UC06 | Zinc finger protein 70 | fenebrutinib |
| Q9BPX5 | Actin-related protein 2/3 complex subunit 5-like protein | Q-203 |
| Q8TDF6 | RAS guanyl-releasing protein 4 | NMS-1286937 |
| Q9P2G3 | Kelch-like protein 14 | NMS-1286937 |
| Q9HBT7 | Zinc finger protein 287 | CCT137690 |
| Q9UKR8 | Tetraspanin-16 | PF-05190457 |
| P59044 | NACHT, LRR and PYD domains-containing protein 6 | MK-5046 |
| Q16774 | Guanylate kinase | MK-5046 |
| Q9HCE5 | N6-adenosine-methyltransferase non-catalytic subunit | PF-05190457 |
| Q15185 | Prostaglandin E synthase 3 | cilofexor |
| Q8WUA7 | TBC1 domain family member 22A | fenebrutinib |
| Q17RS7 | Flap endonuclease GEN homolog 1 | CGM097 |
| Q14244 | Ensconsin | PF-05190457 |
| Q9BRK5 | 45 kDa calcium-binding protein | PF-05190457 |
| Q9UKJ5 | Cysteine-rich hydrophobic domain-containing protein 2 | CCT137690 |
| Q6ZWJ1 | Syntaxin-binding protein 4 | AI-10-49 |
| Q9HAR2 | Adhesion G protein-coupled receptor L3 | AI-10-49 |
| P98161 | Polycystin-1 | AI-10-49 |
| Q92673 | Sortilin-related receptor | AI-10-49 |
| D6RGH6 | Multicilin | fenebrutinib |
| Q8NHH9 | Atlastin-2 | fenebrutinib |
| Q9H0E2 | Toll-interacting protein | PF-05190457 |
| O15145 | Actin-related protein 2/3 complex subunit 3 | PF-05190457 |
| P51116 | Fragile X mental retardation syndrome-related protein 2 | abemaciclib |

**Table 3 continued from previous page**

| Uniprot | Protein name | Drug name |
|---|---|---|
| Q9BR09 | Neuralized-like protein 2 | elbasvir |
| P42568 | Protein AF-9 | AI-10-49 |
| P17600 | Synapsin-1 | AI-10-49 |
| P48553 | Trafficking protein particle complex subunit 10 | AI-10-49 |
| Q12955 | Ankyrin-3 | abemaciclib |
| O60936 | Nucleolar protein 3 | abemaciclib |
| Q02575 | Helix-loop-helix protein 1 | AI-10-49 |
| P49640 | Homeobox even-skipped homolog protein 1 | CFI-402257 |
| P22670 | MHC class II regulatory factor RFX1 | PF-05190457 |
| Q8IUF8 | Ribosomal oxygenase 2 | NMS-1286937 |
| P22681 | E3 ubiquitin-protein ligase CBL | NMS-1286937 |
| Q7RTS3 | Pancreas transcription factor 1 subunit alpha | MK-5046 |
| Q9Y5L4 | Mitochondrial import inner membrane translocase subunit Tim13 | NMS-P715 |
| P17735 | Tyrosine aminotransferase | PF-05190457 |
| O95294 | RasGAP-activating-like protein 1 | PF-05190457 |
| Q8NCD3 | Holliday junction recognition protein | PF-05190457 |
| Q86W28 | NACHT, LRR and PYD domains-containing protein 8 | MK-5046 |
| Q04671 | P protein | AI-10-49 |
| Q9C035 | Tripartite motif-containing protein 5 | AI-10-49 |
| Q66K64 | DDB1- and CUL4-associated factor 15 | AI-10-49 |
| Q5T9A4 | ATPase family AAA domain-containing protein 3B | CFI-402257 |
| Q96MP8 | BTB/POZ domain-containing protein KCTD7 | CFI-402257 |
| Q9UIE0 | Zinc finger protein 230 | CFI-402257 |
| Q3SYG4 | Protein PTHB1 | abemaciclib |
| Q8IUY3 | GRAM domain-containing protein 2A | AI-10-49 |
| O75689 | Arf-GAP with dual PH domain-containing protein 1 | AI-10-49 |
| Q9NZP8 | Complement C1r subcomponent-like protein | fenebrutinib |
| Q9NQX4 | Unconventional myosin-Vc | fenebrutinib |
| Q15388 | Mitochondrial import receptor subunit TOM20 homolog | CFI-402257 |
| Q9NXL9 | DNA helicase MCM9 | piperaquine-phosphate |
| P53370 | Nucleoside diphosphate-linked moiety X motif 6 | piperaquine-phosphate |
| Q8N7C0 | Leucine-rich repeat-containing protein 52 | PF-05190457 |
| O95231 | Homeobox protein VENTX | NMS-P715 |

**Table 3 continued from previous page**

| Uniprot | Protein name | Drug name |
|---|---|---|
| Q9HC56 | Protocadherin-9 | AI-10-49 |
| A6NNW6 | Enolase 4 | AI-10-49 |
| P35612 | Beta-adducin | CFI-402257 |
| Q9Y6M5 | Zinc transporter 1 | AI-10-49 |
| Q01804 | OTU domain-containing protein 4 | AI-10-49 |
| Q9Y678 | Coatomer subunit gamma-1 | MK-5046 |
| Q9BTY7 | Protein HGH1 homolog | AI-10-49 |
| Q8NEM0 | Microcephalin | CFI-402257 |
| Q7LGA3 | Heparan sulfate 2-O-sulfotransferase 1 | AI-10-49 |
| O75956 | Cyclin-dependent kinase 2-associated protein 2 | ziritaxestat |
| P62987 | Ubiquitin-60S ribosomal protein L40 | ziritaxestat |
| Q14257 | Reticulocalbin-2 | CDK9-IN-6 |
| Q9H0I3 | Coiled-coil domain-containing protein 113 | tropifexor |
| P32019 | Type II inositol 1,4,5-trisphosphate 5-phosphatase | fenebrutinib |
| O95409 | Zinc finger protein ZIC 2 | fenebrutinib |
| P56179 | Homeobox protein DLX-6 | fenebrutinib |
| Q12798 | Centrin-1 | AI-10-49 |
| Q14714 | Sarcospan | AI-10-49 |
| Q96LI5 | CCR4-NOT transcription complex subunit 6-like | abemaciclib |
| Q86XR7 | TIR domain-containing adapter molecule 2 | AI-10-49 |
| P53672 | Beta-crystallin A2 | CCT137690 |
| O43439 | Protein CBFA2T2 | CCT137690 |
| Q9H0E3 | Histone deacetylase complex subunit SAP130 | fenebrutinib |
| Q8NFZ0 | F-box DNA helicase 1 | fenebrutinib |
| Q8N0S2 | Synaptonemal complex central element protein 1 | CFI-402257 |
| Q92667 | A-kinase anchor protein 1, mitochondrial | fenebrutinib |
| O60884 | DnaJ homolog subfamily A member 2 | PF-05190457 |
| P41214 | Eukaryotic translation initiation factor 2D | AI-10-49 |
| Q6R327 | Rapamycin-insensitive companion of mTOR | AI-10-49 |
| O75886 | Signal transducing adapter molecule 2 | PF-05190457 |
| Q9UMX6 | Guanylyl cyclase-activating protein 2 | PF-05190457 |
| Q99447 | Ethanolamine-phosphate cytidylyltransferase | CCT137690 |
| Q6FI13 | Histone H2A type 2-A | PF-05190457 |
| Q969S2 | Endonuclease 8-like 2 | fenebrutinib |
| Q6XZF7 | Dynamin-binding protein | AI-10-49 |
| Q96EX1 | Small integral membrane protein 12 | fenebrutinib |

**Table 3 continued from previous page**

| Uniprot | Protein name | Drug name |
|---|---|---|
| Q8TD57 | Dynein axonemal heavy chain 3 | fenebrutinib |
| O15375 | Monocarboxylate transporter 6 | PF-05190457 |
| Q9H7E2 | Tudor domain-containing protein 3 | NMS-1286937 |
| A6NI73 | Leukocyte immunoglobulin-like receptor subfamily A member 5 | AI-10-49 |
| Q9NZM3 | Intersectin-2 | CDK9-IN-6 |
| O75145 | Liprin-alpha-3 | CDK9-IN-6 |
| Q9NSD9 | Phenylalanine–tRNA ligase beta subunit | abemaciclib |
| O15195 | Villin-like protein | abemaciclib |
| P05538 | HLA class II histocompatibility antigen, DQ beta 2 chain | cilofexor |
| Q8TD57 | Dynein axonemal heavy chain 3 | cilofexor |
| Q9NRD9 | Dual oxidase 1 | abemaciclib |
| Q8WUQ7 | Cactin | AI-10-49 |
| Q9H7C4 | Syncoilin | AI-10-49 |
| Q70EK9 | Ubiquitin carboxyl-terminal hydrolase 51 | PF-05190457 |
| O43837 | Isocitrate dehydrogenase [NAD] subunit beta, mitochondrial | PF-05190457 |
| Q9UKY1 | Zinc fingers and homeoboxes protein 1 | CFI-402257 |
| Q6ZMJ2 | Scavenger receptor class A member 5 | CFI-402257 |
| P61371 | Insulin gene enhancer protein ISL-1 | fenebrutinib |
| Q6NSZ9 | Zinc finger and SCAN domain-containing protein 25 | fenebrutinib |
| Q92754 | Transcription factor AP-2 gamma | IACS-10759 |
| Q5RKV6 | Exosome complex component MTR3 | AI-10-49 |
| Q9BRA0 | N-alpha-acetyltransferase 38, NatC auxiliary subunit | MK-5046 |
| Q9BSF8 | BTB/POZ domain-containing protein 10 | CCT137690 |
| Q6PJ69 | Tripartite motif-containing protein 65 | AI-10-49 |
| Q9NRP0 | Oligosaccharyltransferase complex subunit OSTC | AI-10-49 |
| Q8TBM7 | Transmembrane protein 254 | PF-05190457 |
| Q674X7 | Kazrin | elbasvir |
| Q9Y3C4 | EKC/KEOPS complex subunit TPRKB | abemaciclib |
| P59190 | Ras-related protein Rab-15 | abemaciclib |
| Q9NZM3 | Intersectin-2 | PF-05190457 |
| P05455 | Lupus La protein | AI-10-49 |
| Q96KN3 | Homeobox protein PKNOX2 | abemaciclib |
| P54278 | Mismatch repair endonuclease PMS2 | abemaciclib |
| P49069 | Guided entry of tail-anchored proteins factor CAMLG | fenebrutinib |
| Q8IYT4 | Katanin p60 ATPase-containing subunit A-like 2 | AI-10-49 |
| Q8WVI0 | Small integral membrane protein 4 | AI-10-49 |

**Table 3 continued from previous page**

| Uniprot | Protein name | Drug name |
|---|---|---|
| Q9Y6X6 | Unconventional myosin-XVI | MK-5046 |
| Q8N357 | Solute carrier family 35 member F6 | MK-5046 |
| Q96BP2 | Coiled-coil-helix-coiled-coil-helix domain-containing protein 1 | abemaciclib |
| Q9UJQ4 | Sal-like protein 4 | MK-5046 |
| Q9HCQ5 | Polypeptide N-acetylgalactosaminyltransferase 9 | fenebrutinib |
| Q9P2Q2 | FERM domain-containing protein 4A | fenebrutinib |
| Q7Z7M8 | UDP-GlcNAc:betaGal beta-1,3-N-acetylglucosaminyltransferase 8 | NMS-P715 |
| Q9HCK5 | Protein argonaute-4 | PF-05190457 |
| O95405 | Zinc finger FYVE domain-containing protein 9 | AI-10-49 |
| Q92911 | Sodium/iodide cotransporter | PF-05190457 |
| Q8WWI1 | LIM domain only protein 7 | fenebrutinib |
| Q8N3C7 | CAP-Gly domain-containing linker protein 4 | fenebrutinib |
| Q9UBS8 | E3 ubiquitin-protein ligase RNF14 | PF-05190457 |
| Q9BY66 | Lysine-specific demethylase 5D | PF-05190457 |
| Q9UPM8 | AP-4 complex subunit epsilon-1 | abemaciclib |
| Q9BYE0 | Transcription factor HES-7 | MK-5046 |
| Q9BSD7 | Cancer-related nucleoside-triphosphatase | AI-10-49 |
| Q9NXH9 | tRNA | tropifexor |
| Q13948 | Protein CASP | tropifexor |
| Q9C019 | Tripartite motif-containing protein 15 | abemaciclib |
| Q5TA50 | Ceramide-1-phosphate transfer protein | ABT-702 |
| Q7KZF4 | Staphylococcal nuclease domain-containing protein 1 | ABT-702 |
| P08247 | Synaptophysin | fenebrutinib |
| A4IF30 | Solute carrier family 35 member F4 | AI-10-49 |
| Q9NRA8 | Eukaryotic translation initiation factor 4E transporter | AI-10-49 |
| Q96FF9 | Sororin | acalabrutinib |
| P10073 | Zinc finger and SCAN domain-containing protein 22 | PF-05190457 |
| P0CW19 | LIM and senescent cell antigen-like-containing domain protein 3 | PF-05190457 |
| Q9UPM8 | AP-4 complex subunit epsilon-1 | fenebrutinib |
| P08118 | Beta-microseminoprotein | PF-05190457 |
| Q96P16 | Regulation of nuclear pre-mRNA domain-containing protein 1A | fenebrutinib |
| P62280 | 40S ribosomal protein S11 | piperaquine-phosphate |
| Q8TE49 | OTU domain-containing protein 7A | AI-10-49 |

**Table 3 continued from previous page**

| Uniprot | Protein name | Drug name |
|---|---|---|
| Q53HI1 | Protein unc-50 homolog | AI-10-49 |
| Q8WW32 | High mobility group protein B4 | PF-05190457 |
| P57729 | Ras-related protein Rab-38 | PF-05190457 |
| Q8N2M8 | CLK4-associating serine/arginine rich protein | fenebrutinib |
| P25705 | ATP synthase subunit alpha, mitochondrial | fenebrutinib |
| Q9Y5H3 | Protocadherin gamma-A10 | PF-05190457 |
| Q9NU63 | Zinc finger protein 57 homolog | fenebrutinib |
| P43363 | Melanoma-associated antigen 10 | AI-10-49 |
| Q13469 | Nuclear factor of activated T-cells, cytoplasmic 2 | AI-10-49 |
| Q8NG77 | Olfactory receptor 2T12 | AI-10-49 |
| Q9Y2G8 | DnaJ homolog subfamily C member 16 | PF-05190457 |
| Q9ULV8 | E3 ubiquitin-protein ligase CBL-C | fenebrutinib |
| Q96EF0 | Myotubularin-related protein 8 | elbasvir |
| Q8TD16 | Protein bicaudal D homolog 2 | abemaciclib |
| P49459 | Ubiquitin-conjugating enzyme E2 A | abemaciclib |
| Q6XZF7 | Dynamin-binding protein | AI-10-49 |
| Q9Y4I1 | Unconventional myosin-Va | AI-10-49 |
| Q96FL9 | Polypeptide N-acetylgalactosaminyltransferase 14 | MK-5046 |
| Q9HD67 | Unconventional myosin-X | MK-5046 |
| Q13214 | Semaphorin-3B | MK-5046 |
| Q53H47 | Histone-lysine N-methyltransferase SETMAR | MK-5046 |
| P46776 | 60S ribosomal protein L27a | AI-10-49 |
| Q15306 | Interferon regulatory factor 4 | AI-10-49 |
| Q2TAA2 | Isoamyl acetate-hydrolyzing esterase 1 homolog | abemaciclib |
| O43324 | Eukaryotic translation elongation factor 1 epsilon-1 | PF-05190457 |
| Q96R27 | Olfactory receptor 2M4 | fenebrutinib |
| Q5VXU1 | Sodium/potassium-transporting ATPase subunit beta-1-interacting protein 2 | fenebrutinib |
| Q5VWQ8 | Disabled homolog 2-interacting protein | fenebrutinib |
| Q9UJF2 | Ras GTPase-activating protein nGAP | elbasvir |
| O75923 | Dysferlin | fenebrutinib |
| O75592 | E3 ubiquitin-protein ligase MYCBP2 | PF-05190457 |
| Q14714 | Sarcospan | CFI-402257 |
| Q12798 | Centrin-1 | CFI-402257 |
| Q9H9Z2 | Protein lin-28 homolog A | fenebrutinib |
| Q8N2A8 | Mitochondrial cardiolipin hydrolase | fenebrutinib |

**Table 3 continued from previous page**

| Uniprot | Protein name | Drug name |
|---|---|---|
| Q5VV41 | Rho guanine nucleotide exchange factor 16 | abemaciclib |
| Q96QT6 | PHD finger protein 12 | fenebrutinib |
| Q13948 | Protein CASP | AI-10-49 |
| Q9NXH9 | tRNA | AI-10-49 |
| Q969E2 | Secretory carrier-associated membrane protein 4 | fenebrutinib |
| P58107 | Epiplakin | fenebrutinib |
| P55036 | 26S proteasome non-ATPase regulatory subunit 4 | NMS-P715 |
| Q11128 | 4-galactosyl-N-acetylglucosaminide 3-alpha-L-fucosyltransferase FUT5 | AI-10-49 |
| Q86W28 | NACHT, LRR and PYD domains-containing protein 8 | AI-10-49 |
| O43566 | Regulator of G-protein signaling 14 | Q-203 |
| Q9BVI0 | PHD finger protein 20 | Q-203 |
| Q93074 | Mediator of RNA polymerase II transcription subunit 12 | PF-05190457 |
| P51530 | DNA replication ATP-dependent helicase/nuclease DNA2 | PF-05190457 |
| Q6NS38 | DNA oxidative demethylase ALKBH2 | AI-10-49 |
| B4DJY2 | Transmembrane protein 233 | AI-10-49 |
| Q8N807 | Protein disulfide-isomerase-like protein of the testis | PF-05190457 |
| P13929 | Beta-enolase | AI-10-49 |
| Q9NRJ7 | Protocadherin beta-16 | AI-10-49 |
| O95751 | Protein LDOC1 | MK-5046 |
| Q92947 | Glutaryl-CoA dehydrogenase, mitochondrial | fenebrutinib |
| Q96J94 | Piwi-like protein 1 | fenebrutinib |
| O15400 | Syntaxin-7 | adapalene |
| O00192 | Armadillo repeat protein deleted in velo-cardio-facial syndrome | adapalene |
| Q17RD7 | Synaptotagmin-16 | CFI-402257 |
| A0PJZ3 | Glucoside xylosyltransferase 2 | MK-5046 |
| Q53GG5 | PDZ and LIM domain protein 3 | PF-05190457 |
| P20929 | Nebulin | PF-05190457 |
| Q5NDL2 | EGF domain-specific O-linked N-acetylglucosamine transferase | abemaciclib |
| Q10570 | Cleavage and polyadenylation specificity factor subunit 1 | fenebrutinib |
| Q53FE4 | Uncharacterized protein C4orf17 | AI-10-49 |
| Q96T21 | Selenocysteine insertion sequence-binding protein 2 | cilofexor |
| P17980 | 26S proteasome regulatory subunit 6A | PF-05190457 |

**Table 3 continued from previous page**

| Uniprot | Protein name | Drug name |
|---|---|---|
| Q8TE77 | Protein phosphatase Slingshot homolog 3 | tropifexor |
| Q96LD4 | E3 ubiquitin-protein ligase TRIM47 | tropifexor |
| P25205 | DNA replication licensing factor MCM3 | fenebrutinib |
| Q9NQC7 | Ubiquitin carboxyl-terminal hydrolase CYLD | fenebrutinib |
| Q71F23 | Centromere protein U | NMS-P715 |
| P46778 | 60S ribosomal protein L21 | fenebrutinib |
| Q04727 | Transducin-like enhancer protein 4 | AI-10-49 |
| O14556 | Glyceraldehyde-3-phosphate dehydrogenase, testis-specific | AI-10-49 |
| Q92994 | Transcription factor IIIB 90 kDa subunit | fenebrutinib |
| Q15646 | 2'-5'-oligoadenylate synthase-like protein | fenebrutinib |
| O15131 | Importin subunit alpha-6 | fenebrutinib |
| Q14254 | Flotillin-2 | fenebrutinib |
| O75764 | Transcription elongation factor A protein 3 | CFI-402257 |
| P40425 | Pre-B-cell leukemia transcription factor 2 | MK-5046 |
| O43766 | Lipoyl synthase, mitochondrial | PF-05190457 |
| Q969L2 | Protein MAL2 | fenebrutinib |
| Q9NSG2 | Uncharacterized protein C1orf112 | CCT137690 |
| Q9H892 | Tetratricopeptide repeat protein 12 | PF-05190457 |
| O95248 | Myotubularin-related protein 5 | CCT137690 |
| O60481 | Zinc finger protein ZIC 3 | PF-05190457 |
| Q9UBK8 | Methionine synthase reductase | AI-10-49 |
| Q4G0J3 | La-related protein 7 | CCT137690 |
| Q8WW32 | High mobility group protein B4 | CCT137690 |
| P23297 | Protein S100-A1 | MK-5046 |
| Q8TCB7 | tRNA N | elbasvir |
| Q8IXH6 | Tumor protein p53-inducible nuclear protein 2 | abemaciclib |

Table S 4: Predicted ligands for the transcription factors and transcription activity related proteins. * labels the transcription factors.

| Uniprot ID | Protein name | Drug name |
|---|---|---|
| P22670* | MHC class II regulatory factor RFX1 | PF-05190457 |
| P40425* | Pre-B-cell leukemia transcription factor 2 | MK-5046 |
| Q7RTS3* | Pancreas transcription factor 1 subunit alpha | MK-5046 |
| Q13469* | T-cell transcription factor NFAT1 | AI-10-49 |
| Q92754* | Transcription factor AP-2 gamma | IACS-10759 |
| Q92994* | Transcription factor IIIB 90 kDa subunit | Fenebrutinib |
| Q93074* | Mediator of RNA polymerase II transcription subunit 12 | PF-05190457 |
| Q9BVI0* | Transcription factor TZP | Q-203 |
| Q9BYE0* | Transcription factor HES-7 | MK-5046 |
| D6RGH6 | Multicilin | Fenebrutinib |
| O60481 | Zinc finger protein ZIC 3 | PF-05190457 |
| O95231 | VENT-like homeobox protein 2 | NMS-P715 |
| O95409 | Zinc finger protein ZIC 2 | Fenebrutinib |
| P05455 | Sjoegren syndrome type B antigen | AI-10-49 |
| P10073 | Zinc finger and SCAN domain-containing protein 22 | PF-05190457 |
| P42568 | mixed-lineage leukemia translocated to chromosome 3 protein | AI-10-49 |
| P49640 | Homeobox even-skipped homolog protein 1 | CFI-402257 |
| P56179 | Homeobox protein DLX-6 | Fenebrutinib |
| P61371 | Insulin gene enhancer protein ISL-1 | Fenebrutinib |
| Q02575 | Nescient helix loop helix 1 | AI-10-49 |
| Q04727 | Transducin-like enhancer protein 4 | AI-10-49 |
| Q13948 | Protein CASP | Tropifexor |
| Q13948 | Protein CASP | AI-10-49 |
| Q15306 | Interferon regulatory factor 4 | AI-10-49 |
| Q6NSZ9 | Zinc finger and SCAN domain-containing protein 25 | Fenebrutinib |
| Q8WW32 | High mobility group protein B4 | PF-05190457 |
| Q8WW32 | High mobility group protein B4 | CCT137690 |
| Q96KN3 | Homeobox protein PKNOX2 | Abemaciclib |
| Q96QT6 | PHD finger protein 12 | Fenebrutinib |
| Q9C019 | Tripartite motif-containing protein 15 | Abemaciclib |
| Q9C035 | Tripartite motif-containing protein 5 | AI-10-49 |
| Q9H7E2 | Tudor domain-containing protein 3 | NMS-1286937 |
| Q9HBT7 | Zinc finger protein 287 | CCT137690 |
| Q9NU63 | Zinc finger protein 57 homolog | Fenebrutinib |
| Q9UBS8 | E3 ubiquitin-protein ligase RNF14 | PF-05190457 |
| Q9UC06 | Zinc finger protein 70 | Fenebrutinib |
| Q9UIE0 | Zinc finger protein 230 | CFI-402257 |
| Q9UJQ4 | Sal-like protein 4 | MK-5046 |
| Q9UKY1 | Zinc fingers and homeoboxes protein 1 | CFI-402257 |

Table S 5: Chemicals interacted with undruggable human proteins

| Drug name | Clinical phase | Mechanism of Action | Number of targeted proteins |
|---|---|---|---|
| AI-10-49 | Preclinical | core binding factor inhibitor | 63 |
| Fenebrutinib | Phase 2 | Bruton's tyrosine kinase (BTK) inhibitor | 56 |
| PF-05190457 | Phase 2 | growth hormone secretagogue receptor inverse agonist | 42 |
| Abemaciclib | Launched | CDK inhibitor | 21 |
| MK-5046 | Preclinical | bombesin receptor agonist | 18 |
| CFI-402257 | Phase 1/Phase 2 | dual specificity protein kinase inhibitor | 14 |
| CCT137690 | Preclinical | Aurora kinase inhibitor | 11 |
| Tropifexor | Phase 2 | FXR agonist | 5 |
| NMS-1286937 | Phase 2 | PLK inhibitor | 5 |
| NMS-P715 | Preclinical | protein kinase inhibitor | 5 |
| Elbasvir | Launched | HCV inhibitor | 5 |
| Cilofexor | Phase 3 | FXR agonist | 4 |
| CDK9-IN-6 | Preclinical | CDK inhibitor | 3 |
| piperaquine-phosphate | Launched | antimalarial agent | 3 |
| Q-203 | Phase 2 | ATP synthase inhibitor | 3 |
| ABT 702 dihydrochloride | Preclinical | adenosine kinase inhibitor | 2 |
| Ziritaxestat | Phase 3 | autotaxin inhibitor | 2 |
| Adapalene | Launched | retinoid receptor agonist | 1 |
| Acalabrutinib | Launched | Bruton's tyrosine kinase (BTK) inhibitor | 1 |
| IACS-10759 Hydrochloride | Preclinical | mitochondrial complex I inhibitor | 1 |
| NVP-CGM097 | Phase 1 | MDM inhibitor | 1 |

Table S 6: Targets predicted by PortalCG for AI-10-49, fenebrutinib, PF-05190457 and their docking score from Autodock Vina.

| Target protein | Drug name | Autodock Vina docking score (kcal/mol) |
|---|---|---|
| Q92673 | AI-10-49 | -7.3 |
| P42568 | AI-10-49 | -6.8 |
| O75689 | AI-10-49 | -5.6 |
| Q86XR7 | AI-10-49 | -10.1 |
| Q5RKV6 | AI-10-49 | -7.7 |
| Q13469 | AI-10-49 | -8.6 |
| O14556 | AI-10-49 | -9.8 |
| Q9UBK8 | AI-10-49 | -8.2 |
| Q9NQX4 | fenebrutinib | -9.5 |
| Q9ULV8 | fenebrutinib | -7.3 |
| O75923 | fenebrutinib | -8.1 |
| Q9H9Z2 | fenebrutinib | -8.2 |
| Q96J94 | fenebrutinib | -8.2 |
| Q9NQC7 | fenebrutinib | -8.3 |
| Q9UKR8 | PF-05190457 | -8.3 |
| P17735 | PF-05190457 | -7.3 |
| O75886 | PF-05190457 | -8.4 |
| Q9NZM3 | PF-05190457 | -7.6 |
| P08118 | PF-05190457 | -7.9 |
| P57729 | PF-05190457 | -8.6 |
| P51530 | PF-05190457 | -9.4 |
| Q53GG5 | PF-05190457 | -7.7 |

Table S 7: Functional Annotation enrichment for human proteins in Tbio selected by PortalCG

| David Functional Annotation enrichment analysis | | | | |
|---|---|---|---|---|
| Enriched terms in UniProtKB keywords | Number of proteins involved | Percentage of proteins involved | P-value | Modified Benjamini p-value |
| Alternative splicing | 153 | 69.9 | 2.70E-08 | 6.80E-06 |
| Phosphoprotein | 126 | 57.5 | 1.70E-07 | 2.20E-05 |
| Cytoplasm | 82 | 37.4 | 3.20E-06 | 2.40E-04 |
| Nucleus | 87 | 39.7 | 3.70E-06 | 2.40E-04 |
| Metal-binding | 61 | 27.9 | 2.00E-04 | 1.00E-02 |

Table S 8: Chemicals interacted with human proteins in Tbio

| rug name | Clinical phase | Mechanism of Action | Number of targeted proteins |
|---|---|---|---|
| AI-10-49 | Preclinical | core binding factor inhibitor | 52 |
| fenebrutinib | Phase 2 | Bruton's tyrosine kinase (BTK) inhibitor | 45 |
| PF-05190457 | Phase 2 | growth hormone secretagogue receptor inverse agonist | 36 |
| abemaciclib | Launched | CDK inhibitor | 20 |
| MK-5046 | Preclinical | bombesin receptor agonist | 15 |
| CFI-402257 | Phase 1/Phase 2 | dual specificity protein kinase inhibitor | 14 |
| CCT137690 | Preclinical | Aurora kinase inhibitor | 7 |
| NMS-1286937 | Phase 2 | PLK inhibitor | 5 |
| tropifexor | Phase 2 | FXR agonist | 4 |
| NMS-P715 | Preclinical | protein kinase inhibitor | 4 |
| cilofexor | Phase 3 | FXR agonist | 3 |
| CDK9-IN-6 | Preclinical | CDK inhibitor | 3 |
| piperaquine-phosphate | Launched | antimalarial agent | 3 |
| elbasvir | Launched | HCV inhibitor | 3 |
| ABT-702 | Preclinical | adenosine kinase inhibitor | 2 |
| adapalene | Launched | retinoid receptor agonist | 2 |
| Q-203 | Phase 2 | ATP synthase inhibitor | 2 |
| ziritaxestat | Phase 3 | autotaxin inhibitor | 2 |
| acalabrutinib | Launched | Bruton's tyrosine kinase (BTK) inhibitor | 1 |
| IACS-10759 | Preclinical | mitochondrial complex I inhibitor | 1 |
| CGM097 | Phase 1 | MDM inhibitor | 1 |

Table S 9: Functional Annotation enrichment for undruggable human disease proteins without Tbio selected by PortalCG

| David Functional Annotation enrichment analysis | | | | |
|---|---|---|---|---|
| Enriched terms in UniProtKB keyword | Number of proteins involved | Percentage of proteins involved (%) | P-value | Modified Benjamini p-value |
| Zinc-finger | 80 | 23 | 7.60E-16 | 1.20E-13 |
| Zinc | 85 | 24.4 | 1.20E-11 | 9.90E-10 |
| Metal-binding | 96 | 27.6 | 4.30E-06 | 2.30E-04 |
| DNA-binding | 62 | 17.8 | 7.90E-06 | 3.20E-04 |
| Transcription regulation | 61 | 17.5 | 5.60E-04 | 1.80E-02 |
| Transcription | 61 | 17.5 | 1.10E-03 | 3.00E-02 |

Table S 10: Top ranked diseases associated with undruggable human proteins excluding Tbio selected by PortalCG

| Disease Name | # of undruggable proteins associated with the disease |
|---|---|
| Body Height | 31 |
| Colorectal Carcinoma | 28 |
| Malignant neoplasm of breast | 26 |
| Breast Carcinoma | 18 |
| Blood Protein Measurement | 18 |
| Leukemia, Myelocytic, Acute | 17 |
| Carcinogenesis | 17 |
| Neoplasm Metastasis | 17 |
| Liver carcinoma | 15 |
| Malignant neoplasm of prostate | 15 |

Table S 11: Chemicals interacted with undruggable human proteins excluding Tbio

| Drug_name | Clinical phase | Mechanism of Action | Number of targeted proteins |
|---|---|---|---|
| fenebrutinib | Phase 2 | Bruton's tyrosine kinase (BTK) inhibitor | 80 |
| PF-05190457 | Phase 2 | growth hormone secretagogue receptor inverse agonist | 50 |
| MK-5046 | Preclinical | bombesin receptor agonist | 38 |
| CCT137690 | Preclinical | Aurora kinase inhibitor | 36 |
| AI-10-49 | Preclinical | core binding factor inhibitor | 31 |
| abemaciclib | Launched | CDK inhibitor | 26 |
| CFI-402257 | Phase 1/Phase 2 | dual specificity protein kinase inhibitor | 20 |
| NMS-P715 | Preclinical | protein kinase inhibitor | 14 |
| NMS-1286937 | Phase 2 | PLK inhibitor | 11 |
| elbasvir | Launched | HCV inhibitor | 8 |
| cilofexor | Phase 3 | FXR agonist | 7 |
| ABBV-744 | Phase 1 | bromodomain inhibitor | 7 |
| tropifexor | Phase 2 | FXR agonist | 7 |
| CDK9-IN-6 | Preclinical | CDK inhibitor | 6 |
| adapalene | Launched | retinoid receptor agonist | 4 |
| Q-203 | Phase 2 | ATP synthase inhibitor | 4 |
| PLX8394 | Phase 1/Phase 2 | serine/threonine kinase inhibitor | 4 |
| ABT-702 | Preclinical | adenosine kinase inhibitor | 4 |
| NVP-BSK805 | Preclinical | JAK inhibitor | 3 |
| OTS167 | Phase 1/Phase 2 | maternal embryonic leucine zipper kinase inhibitor | 3 |
| CHIR-99021 | Preclinical | glycogen synthase kinase inhibitor | 3 |
| DBPR-211 | Preclinical | cannabinoid receptor antagonist | 2 |
| A-887826 | Preclinical | sodium channel blocker | 2 |
| integrin-antagonist-1 | Phase 1 | integrin antagonist | 2 |
| piperaquine-phosphate | Launched | antimalarial agent | 2 |
| cenerimod | Phase 2 | sphingosine 1-phosphate receptor modulator | 1 |
| peposertib | Phase 1/Phase 2 | DNA dependent protein kinase inhibitor | 1 |
| tezacaftor | Launched | CFTR channel agonist | 1 |
| cot-inhibitor-2 | Preclinical | MAPK-interacting kinase inhibitor | 1 |
| itacitinib | Phase 3 | JAK inhibitor | 1 |
| 10-hydroxycamptothecin | Preclinical | topoisomerase inhibitor | 1 |
| alectinib | Launched | ALK tyrosine kinase receptor inhibitor | 1 |
| adarotene | Phase 1 | retinoid receptor agonist | 1 |
| acalabrutinib | Launched | Bruton's tyrosine kinase (BTK) inhibitor | 1 |
| XL041 | Preclinical | LXR agonist | 1 |
| WAY-207024 | Preclinical | gonadotropin releasing factor hormone receptor antagonist | 1 |
| MK-5108 | Phase 1 | Aurora kinase inhibitor | 1 |
| CGM097 | Phase 1 | MDM inhibitor | 1 |
| CD-437 | Preclinical | retinoid receptor agonist | 1 |
| AMG-925 | Phase 1 | CDK inhibitor\|FLT3 inhibitor | 1 |
| ACT-132577 | Launched | endothelin receptor antagonist | 1 |
| ziritaxestat | Phase 3 | autotaxin inhibitor | 1 |

### 3 Additional figures

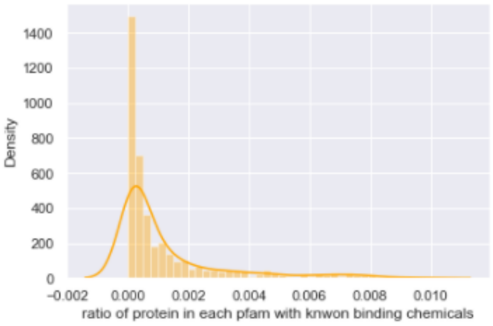

Figure S 1: Dark space statistics histogram based on known CPI pairs in ChEMBL26. < 1% proteins in each pfam invovled in ChEMBLE have known binding chemcials.

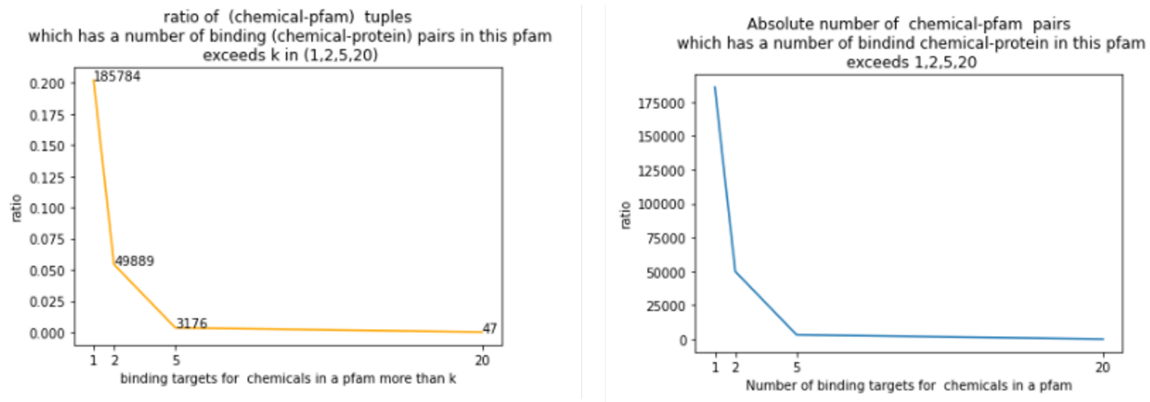

Figure S 2: Dark space statistics histogram based on known CPI pairs in ChEMBL26. < 1% chemicals bind to more then 2 proteins; < 0.4% chemicals bind to more than 5 proteins.

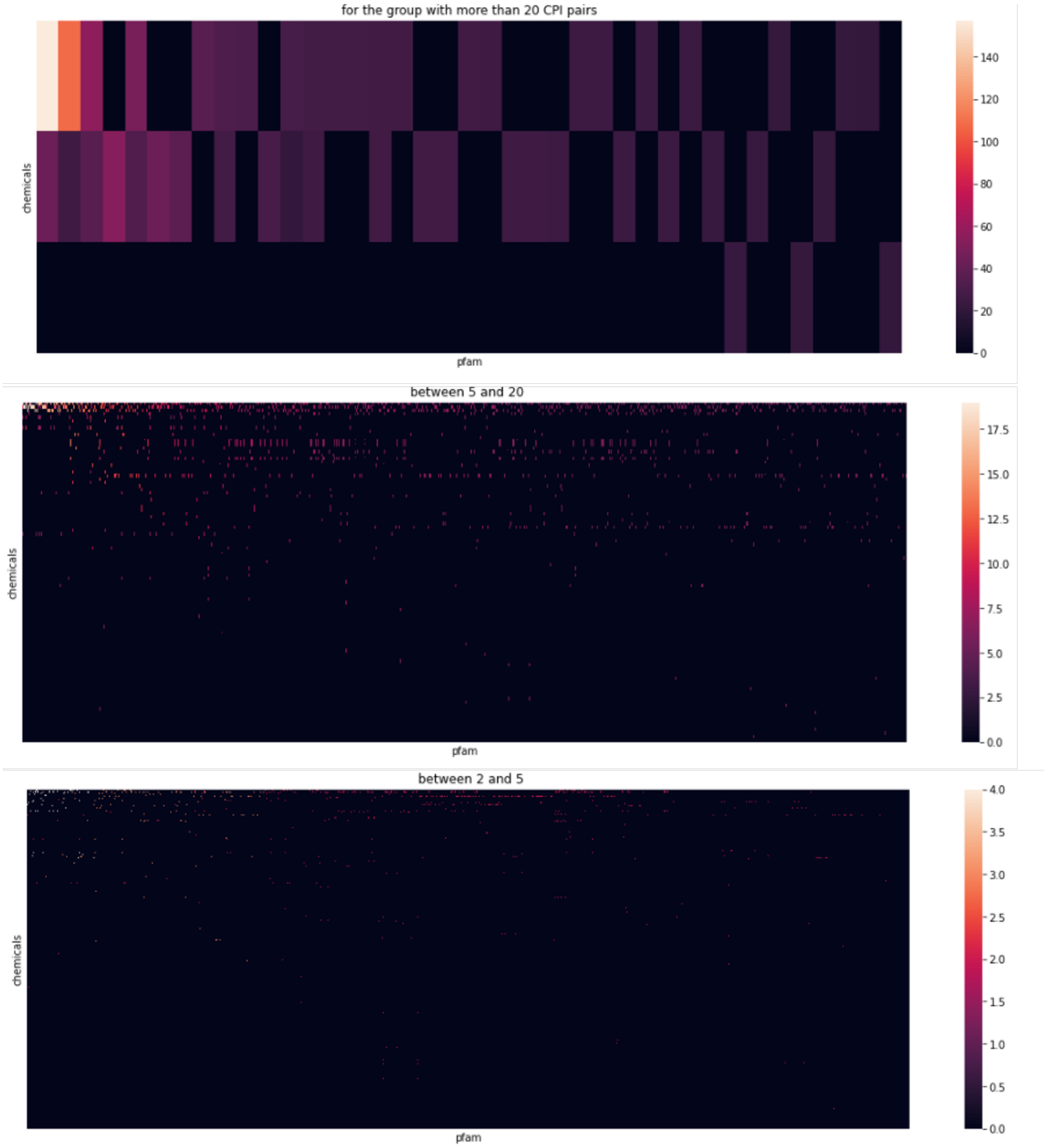

Figure S 3: As shown in Figure S 2, there are three main ranges in terms of the binding targets in a pfam for one chemical: [2,5],[5,20],[20,). For each of the range, a heatmap is shown with y axis representing each chemicals, x axis representing each pfam, each point representing the known binding pairs for one chemical and one pfam. As we can see, there is huge dark space.

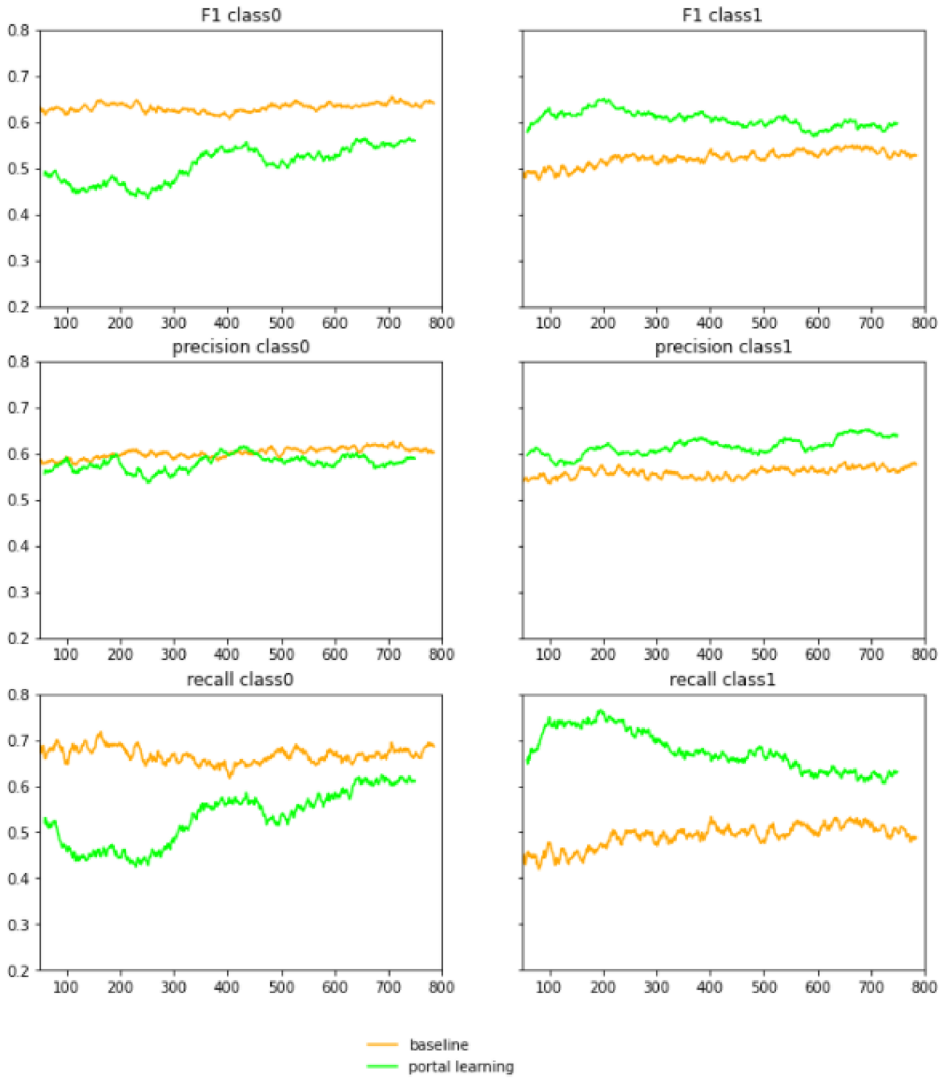

Figure S 4: In the main text, overall evaluation across positive and negative classes are reported, such as F1, ROC-AUC, PR-AUC. Here is a breakdown of performance in each class, where class0 is negative, i.e. not binding, class1 is positive, i.e. binding. against DISAE as baseline

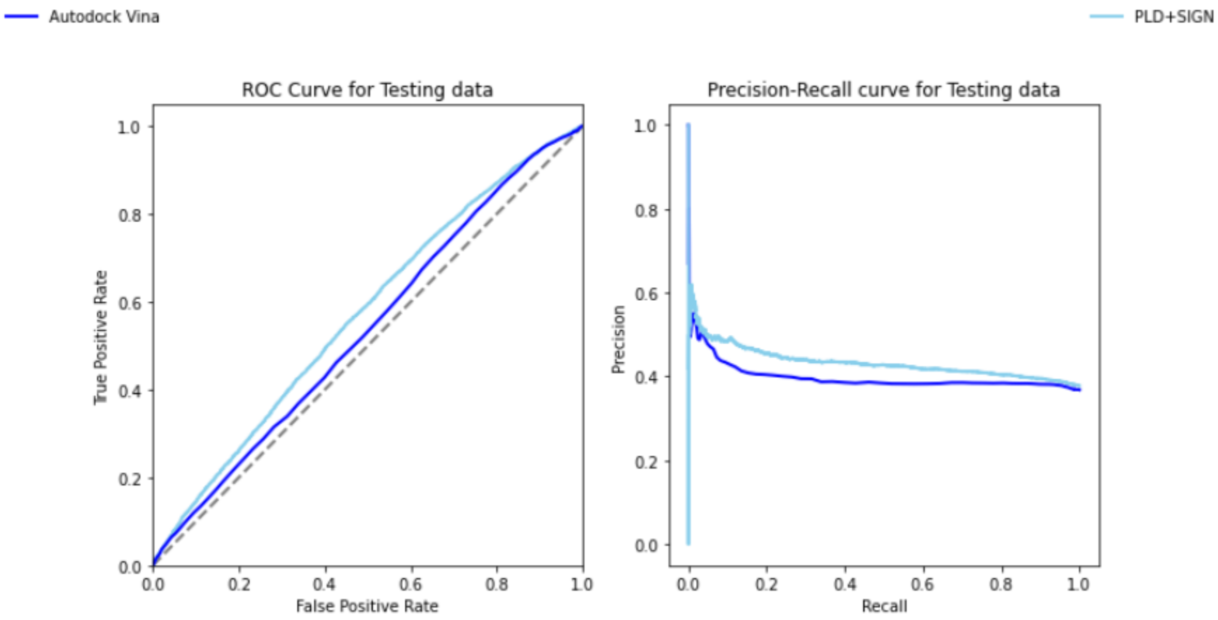

Figure S 5: Performance comparison of PLD+SIGN and Autodock Vina using the same OOD-test set as in main text Figure 3: ROC-AUCs of Autodock Vina and PLD+SIGN are 0.535 and 0.569, respectively. PR-AUCs of Autodock Vina and PLD+SIGN 0.398 and 0.433, respectively.

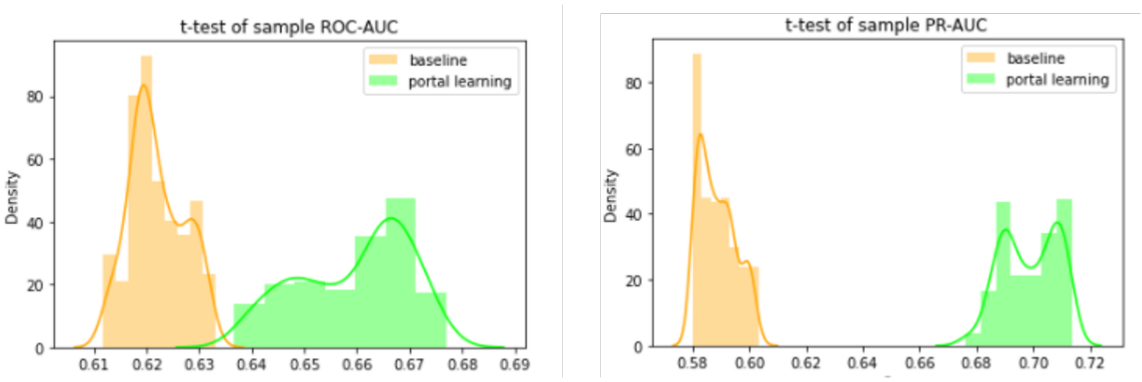

Figure S 6: t-test comparison. The p-values for both ROC-AUC and PR-AUC are close to 0 against DISAE as baseline

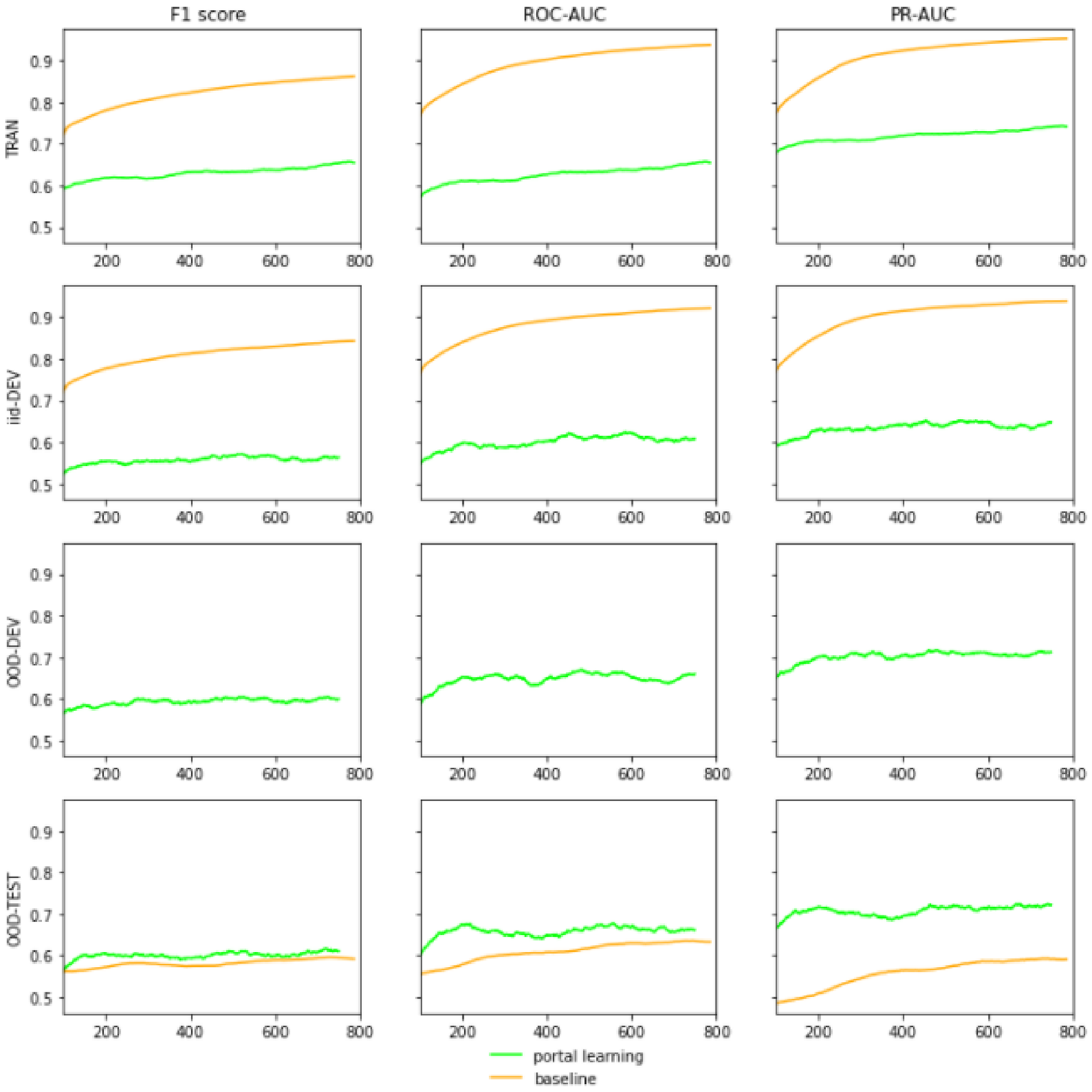

Figure S 7: Stress model selection performance curves against DISAE as baseline

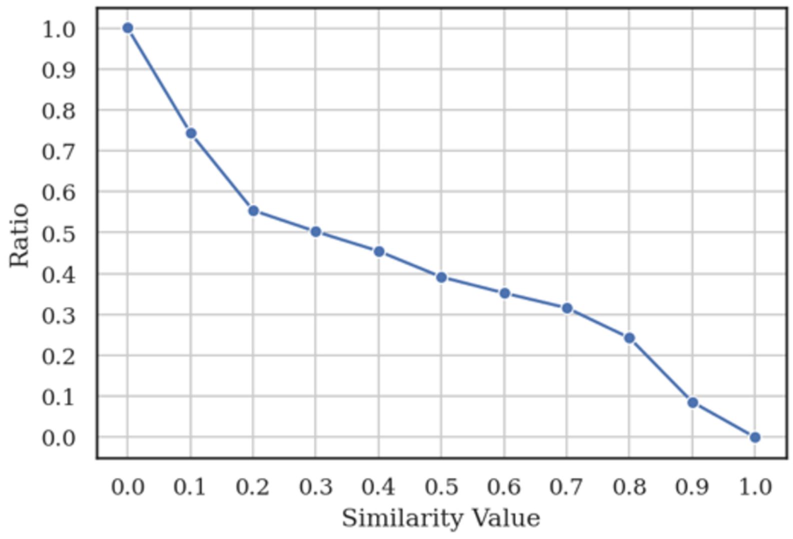

Figure S 8: The ratio of inactive CPIs vs active CPIs under different Tanimoto coefficients of chemical similarities in the training data. The ratio of total inactive CPIs vs active CPIs is 1:1.

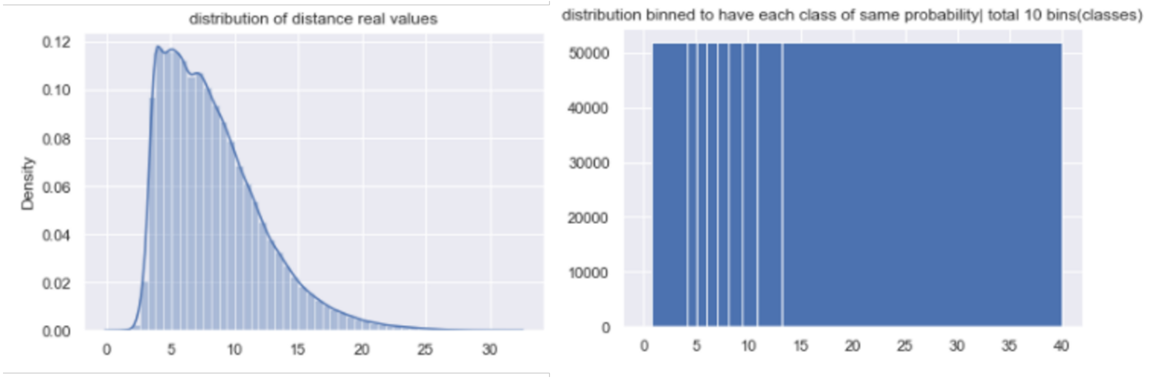

Figure S 9: Histogram equalization results: the left panel shows the original distribution of distance real values; to formalize a multi-class classification where each class has equal probability, histogram equalization transforms the distribution to the right panel of 10 bins, each as a class.

### References

[1] O. Trott and A. J. Olson, “Autodock vina: improving the speed and accuracy of docking with a new scoring function, efficient optimization and multithreading,” *Journal of Computational Chemistry*, vol. 31, pp. 455–461, 2010.

[2] S. Madapa, S. Gadhiya, T. Kurtzman, I. L. Alberts, S. Ramsey, M. Reith, and W. W. Harding, “Synthesis and evaluation of c9 alkoxy analogues of (-)-stepholidine as dopamine receptor ligands,” *European journal of medicinal chemistry*, vol. 125, pp. 255–268, 2017.

[3] R. K. Pal, S. Gadhiya, S. Ramsey, P. Cordone, L. Wickstrom, W. W. Harding, T. Kurtzman, and E. Gallicchio, “Inclusion of enclosed hydration effects in the binding free energy estimation of dopamine d3 receptor complexes,” *Plos one*, vol. 14, no. 9, p. e0222902, 2019.

[4] S. Gadhiya, P. Cordone, R. K. Pal, E. Gallicchio, L. Wickstrom, T. Kurtzman, S. Ramsey, and W. W. Harding, “New dopamine d3-selective receptor ligands containing a 6-methoxy-1, 2, 3, 4-tetrahydroisoquinolin-7-ol motif,” *ACS medicinal chemistry letters*, vol. 9, no. 10, pp. 990–995, 2018.

[5] P. Cordone, H. K. Namballa, B. Muniz, R. K. Pal, E. Gallicchio, and W. W. Harding, “New tetrahydroisoquinoline-based d3r ligands with an o-xylenyl linker motif,” *Bioorganic & Medicinal Chemistry Letters*, vol. 42, p. 128047, 2021.
